# AFLAP: Assembly-Free Linkage Analysis Pipeline using *k*-mers from whole genome sequencing data

**DOI:** 10.1101/2020.09.14.296525

**Authors:** Kyle Fletcher, Lin Zhang, Juliana Gil, Rongkui Han, Keri Cavanaugh, Richard Michelmore

## Abstract

**Background:** Genetic maps are an important resource for validation of genome assemblies, trait discovery, and breeding. Next generation sequencing has enabled production of high-density genetic maps constructed with 10,000s of markers. Most current approaches require a genome assembly to identify markers. Our Assembly Free Linkage Analysis Pipeline (AFLAP) removes this requirement by using uniquely segregating *k*-mers as markers to rapidly construct a genotype table and perform subsequent linkage analysis. This avoids potential biases including preferential read alignment and variant calling.

**Results:** The performance of AFLAP was determined in simulations and contrasted to a conventional workflow. We tested AFLAP using 100 F_2_ individuals of *Arabidopsis thaliana*, sequenced to low coverage. Genetic maps generated using *k*-mers contained over 130,000 markers that were concordant with the genomic assembly. The utility of AFLAP was then demonstrated by generating an accurate genetic map using genotyping-by-sequencing data of 235 recombinant inbred lines of *Lactuca* spp. AFLAP was then applied to 83 F_1_ individuals of the oomycete *Bremia lactucae*, sequenced to >5x coverage. The genetic map contained over 90,000 markers ordered in 19 large linkage groups. This genetic map was used to fragment, order, orient, and scaffold the genome, resulting in a much-improved reference assembly.

**Conclusions:** AFLAP can be used to generate high density linkage maps and improve genome assemblies of any organism when a mapping population is available using whole genome sequencing or genotyping-by-sequencing data. Genetic maps produced for *B. lactucae* were accurately aligned to the genome and guided significant improvements of the reference assembly.

## Background

The complexity of contemporary maps has increased drastically since linkage, the tendency for co-segregation of two or more genetic loci during meiosis relative to their proximity on a chromosome, was introduced as a concept at the start of the 20^th^ century (1, 2). The first genetic map was based on six sex-linked phenotypes in *Drosophila melanogaster* (3). Now, due to technical advances, particularly in DNA sequencing, it is common to construct genetic maps with thousands of markers that greatly exceed the number of genetic bins observed in segregating progeny. Typically, markers are derived from aligning sequencing reads to a reference assembly and calling polymorphisms. However, high quality reference assemblies are not available for all species and may not be available for all genotypes, even in well-studied species. Therefore, we developed a pipeline for generating ultra-high-density genetic maps that does not require a genome assembly and can be applied to any species regardless of genomic resources.

*Bremia lactucae* is an outbreeding oomycete that causes the economically important downy mildew disease of lettuce. Multiple linkage studies have achieved increasing marker density over time. The first genetic map for *B. lactucae*, reporting 13 linkage groups, was generated using 53 restriction fragment length polymorphisms (RFLPs) and nine phenotypic loci spanning 230 cM (4). A second map was based on 83 RFLPs, 347 amplified fragment length polymorphisms (AFLPs), and seven phenotypic loci (5). Genomic investigations of *B. lactucae* revealed genetic characteristics that complicated map construction. Although *B. lactucae* was shown to be a diploid species, many isolates were determined to be heterokaryotic with genetically distinct, diploid nuclei sharing the same coenocytic cytoplasm (6). Single nucleotide polymorphism (SNP) analysis of the F_1_ population previously used for map construction revealed two groups of half-siblings, indicating that one of the parents was contributing two sets of gametes. In addition, analysis of multiple isolates revealed high levels of heterozygosity (> 1%) and the reference assembly consisted of over 70% long terminal repeat retrotransposons (6). These genomic features may reduce the accuracy of short-read mapping and SNP calling (7), which are critical for genetic map construction with next generation sequencing data. Therefore, at the beginning of this study, the genetic architecture of *B. lactucae* remained unresolved.

Our Assembly Free Linkage Analysis Pipeline (AFLAP) enables the construction of genetic maps without mapping or SNP calling against a reference genome assembly. Instead, reads are reduced to *k*-mers (*k*=31, onward referred to as 31-mer) and surveyed to identify those that segregate uniquely in the gametes of each parent. AFLAP was benchmarked against a conventional linkage analysis pipeline. Simulations were used to investigate the impacts of varying genome size, heterozygosity and sequencing depth on run times and results produced by AFLAP. Running AFLAP on an F_2_ population of *Arabidopsis thaliana* Colombia (Col) x Landsberg (Ler) whole genome sequenced to low coverage generated the five expected linkage groups. Testing AFLAP on reduced representation genotype-by-sequencing (GBS) data of a recombinant inbred line population of a *Lactuca serriola* x *L. sativa* interspecific-cross generated the expected nine linkage groups for both parents. AFLAP was then used to construct an ultra-dense genetic map for *B. lactucae*, using F_1_ isolates that had been sequenced to over 5x by whole genome sequencing (WGS).

## Results

A novel Assembly Free Linkage Analysis Pipeline (AFLAP) was designed to rapidly construct genetic maps without requiring the alignment of reads to, and subsequent variant calling against a reference genome assembly (Figure 1). Briefly, AFLAP generated *k*-mers of a set length (here, 31 base pairs referred onward as 31-mer) from the parental read sets. Single copy 31-mers were identified by analyzing peaks contained within count files. Single copy 31-mers common to both parents were discarded; 31-mers unique to each parent were assembled creating variants unique to either parent. One 31-mer was extracted from each fragment of 61 bp and larger to be used as markers. These markers were then surveyed in 31-mers present in progeny individuals. Genotypes were scored as present or absent, resulting in a genotype table, which was then inspected to obtain segregation statistics of the markers. Markers were filtered for segregation distortion and finally exported to LepMap3 (8) for linkage analysis. The pipeline is described in detail in the Materials and Methods section.

**Figure 1.**
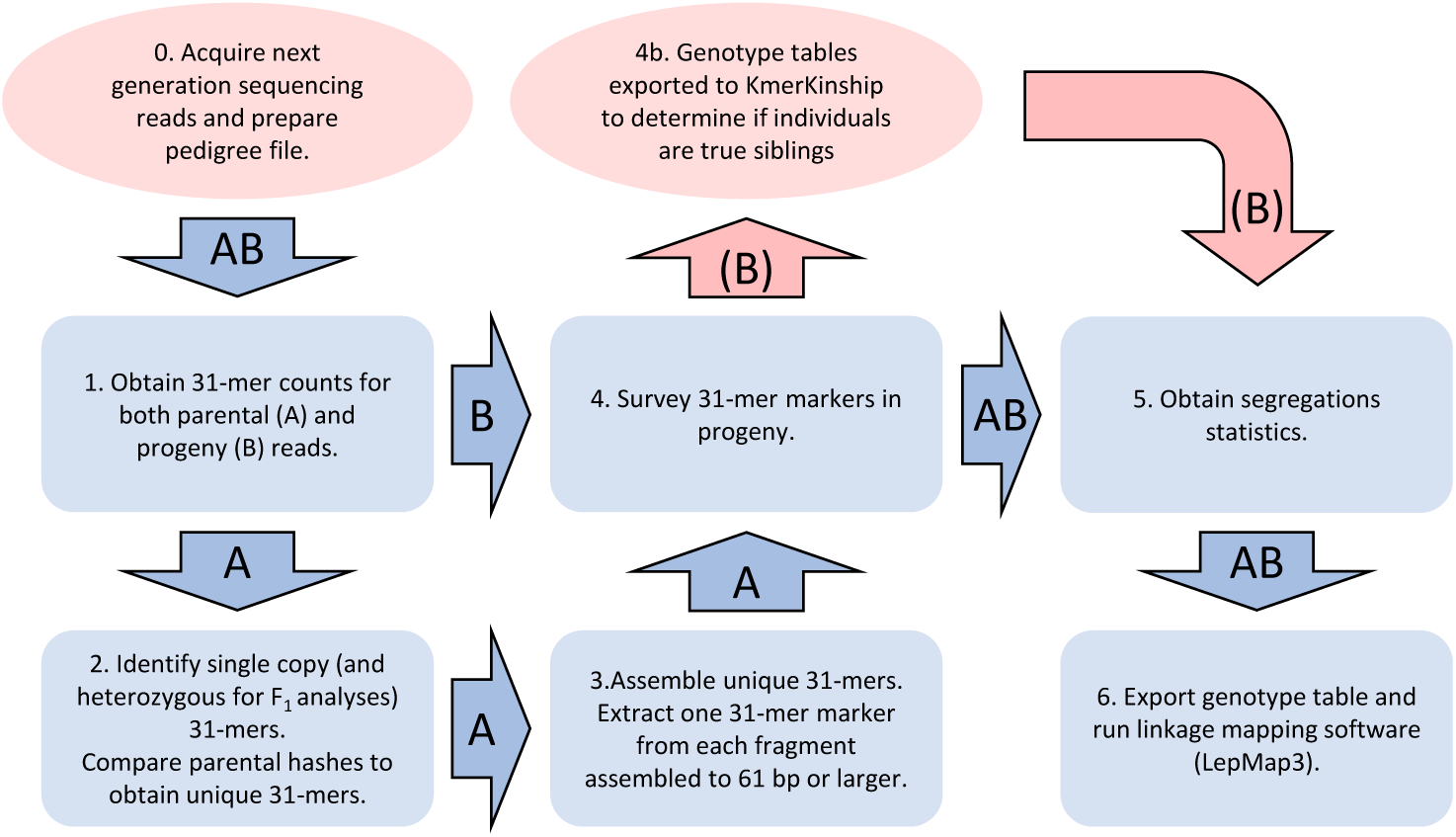
Assembly Free Linkage Analysis Pipeline (AFLAP). Blue rectangles summarize core steps of the pipeline and numbers correspond to script numbering found at https://github.com/kfletcher88/AFLAP. Arrows indicate where one step outputs the input of the next step. Arrows labelled A indicate the path taken in genotyping and identifying markers from parental individuals. Arrows labelled B indicate the path taken to genotype progeny individuals. Red ovals and arrows are supplementary to AFLAP and not required to complete the pipeline. An example pedigree file is provided for the *A. thaliana* data (Additional File 2: Table S1).

### Simulations

To benchmark AFLAP, we simulated a test-cross population of 100 F_1_ individuals and compared the results from AFLAP to those from a conventional pipeline. The 119 Mb, five-chromosome *Arabidopsis thaliana* genome assembly was used as a template to simulate one parent to be 0.2% heterozygous with ∼ 118,000 SNPs and ∼ 1,200 indels. One hundred F_1_ progeny were simulated by introducing one or two crossovers per chromosome (see Materials and Methods). Running AFLAP on synthetic reads derived from the simulated F_1_ progeny resulted in a 699 cM genetic map containing five linkage groups. The average Kendall Rank Coefficient (τ) per linkage group was 0.986, demonstrating that the results were colinear with the reference assembly (Table 1, Additional File 1). This dataset was compared to a conventional read mapping, variant calling, and linkage analysis pipeline, with the same linkage analysis software. The conventional pipeline produced similar results, calculating a 701 cM genetic maps containing five linkage groups that correlated with the reference assembly (τ = 0.992; Table 1, Additional File 1). Significantly, AFLAP was approximately three times as fast as the conventional pipeline and could be further accelerated by down-sampling markers. The average τ across linkage groups showed that down-sampling markers did not alter the correlation of the genetic map with the reference assembly, nor did the map length or number of linkage groups change (Table 1). Therefore, AFLAP can produce accurate genotype tables for linkage analysis much faster without using a genome assembly and results are comparable to conventional pipelines.

**Table 1.** Comparison of AFLAP with a conventional linkage analysis pipeline.

| Method | Assembly size | Chromosomes | Segregating Markers | Markers Used | Minimum LOD score | Linkage Groups | Map Length | Percent of Markers placed | Average Kendall Rank Coefficient ($\tau$) | Run Time |
| --- | --- | --- | --- | --- | --- | --- | --- | --- | --- | --- |
| AFLAP | 119 Mb | 5 | 186,523 | 186,523 | 7 | 5 | 699 cM | 99.8% | 0.986 | 59 hrs 32 mins |
| AFLAP | 119 Mb | 5 | 186,523 | 10,000 | 7 | 5 | 696 cM | 99.8% | 0.986 | 6 hrs 13 mins |
| Conventional | 119 Mb | 5 | 225,797 | 225,797 | 20 | 5 | 701 cM | 99.6% | 0.992 | 173 hrs 34 mins |
| Conventional | 119 Mb | 5 | 226,087 | 10,000 | 7 | 5 | 698 cM | 99.8% | 0.992 | 107 hrs 5 mins |

Performance of AFLAP was investigated by simulating different biological and experimental inputs into the pipeline. The most significant factor impacting AFLAP was the sequencing depth of the progeny (Table 2, Additional File 1). When progenies were simulated to have adequate sequence coverage (≥5x), the linkage groups were highly colinear with the reference assembly with ≥98% of markers placed and average τ > 0.98 across linkage groups. When the simulated sequencing depth was reduced to 3x, only 86.7% of the markers were placed in linkage groups and marker order was less correlated with the reference assembly (τ = 0.826; Table 2, Additional File 1). The sequencing coverage of the parents had less of an effect on the final map quality, with maps being colinear (τ > 0.98); however, the number of markers reported reduced as the coverage dropped (Table 2). Therefore, AFLAP can use low coverage (10x) parental sequencing data to produce accurate genetic maps, albeit with potentially lower information content. The effect of heterozygosity was tested by varying the synthetic heterozygosity (from 0.001% to 2%) of the mapped parent. Genetic maps produced were colinear with the original assembly (τ > 0.98), suggesting that heterozygosity had little effect on the calling of markers (Table 2, Additional File 1). Through down-sampling markers to 10,000, AFLAP was able to construct genetic maps from all simulations with the 119 Mb, five-chromosome genome in under nine hours under the test conditions (Table 2). The impact of different genome sizes and chromosome numbers was tested by simulating crosses using the genomes of *Vitis riparia* (19 chromosomes, ∼500 Mb reference assembly) and *Atriplex hortensis* (nine chromosomes, ∼900 Mb reference assembly), synthesizing the mapped parent to be 0.2% heterozygous and sequencing coverage to be 50x for the parents, 10x for the progeny making it directly comparable with previous simulations. As expected, the number of reported markers increased with genome size. As with the smaller 119 Mb genome, the genetic maps produced with down-sampled markers were colinear with genome assembly from which they were derived (τ > 0.98; Table 2, Additional File 1); however, more markers were required after down-sampling. Fifty thousand markers produced a synthetic genetic map colinear with the 19-chromosome, 500 Mb reference; simulations with 10,000 and 25,000 markers resulted in at least one error. For the largest genome (900 Mb), 25,000 markers were able to reconstitute the nine chromosomes (Table 2, Additional File 1), suggesting that the number of chromosomes, not the genome size is the driver for requiring more markers in these simulations. The time required to run AFLAP increased with genome size, although it was still faster than the conventional pipeline on the smallest simulated cross (Table 1, 2). These simulations demonstrated that AFLAP can accommodate genomes of differing sizes, chromosome numbers, and heterozygosity provided adequate sequencing depth of the progeny is available.

Finally, AFLAP was tested on 100 synthetic F_2_ individuals. For this simulation, the five-chromosome, 119 Mb genomes of both parents were 100% homozygous, varying from one another by ∼ 118,000 SNPs and ∼ 1,200 indels. Whole genome sequencing coverage was simulated to 50x for both parents and 10x for the F_2_ progeny. The resultant five linkage groups in the genetic maps of each parent were colinear with the reference assembly (τ > 0.97; Table 2, Additional File 1). Therefore, AFLAP can be applied to F_1_ and F_2_ populations, and could be extended to other types of populations such as recombinant inbred lines.

**Table 2.** Simulation of AFLAP testing different genome and sequencing coverage variations.

| Assembly size | Chromosomes | Cross structure | Progeny sequencing depth | Parental sequencing depth | Heterozygosity | Segregating Markers | Markers Used | Minimum LOD score | Linkage Groups | Percent of Markers placed | Average Kendall Rank Coefficient ($\tau$) | Run Time |
| --- | --- | --- | --- | --- | --- | --- | --- | --- | --- | --- | --- | --- |
| i) Altering progeny sequencing depth. |  |  |  |  |  |  |  |  |  |  |  |  |
| 119 Mb | 5 | F<sub>1</sub> (ABxCC) | 20 | 50 | 0.2% | 186,522 | 10,000 | 7 | 5 | 100% | 0.993 | 8 hrs 44 mins |
| 119 Mb | 5 | F<sub>1</sub> (ABxCC) | 10 | 50 | 0.2% | 186,523 | 10,000 | 7 | 5 | 99.8% | 0.986 | 6 hrs 13 mins |
| 119 Mb | 5 | F<sub>1</sub> (ABxCC) | 7 | 50 | 0.2% | 186,523 | 10,000 | 7 | 5 | >99.9% | 0.988 | 5 hrs 18 mins |
| 119 Mb | 5 | F<sub>1</sub> (ABxCC) | 5 | 50 | 0.2% | 186,525 | 10,000 | 7 | 5 | 98% | 0.983 | 4 hrs 22 mins |
| 119 Mb | 5 | F<sub>1</sub> (ABxCC) | 3 | 50 | 0.2% | 186,525 | 10,000 | 3 | 6 | 86.7% | 0.826 | 4 hrs 17 mins |
| ii) Altering parental sequencing depth. |  |  |  |  |  |  |  |  |  |  |  |  |
| 119 Mb | 5 | F<sub>1</sub> (ABxCC) | 10 | 50 | 0.2% | 186,523 | 10,000 | 7 | 5 | 99.8% | 0.986 | 6 hrs 13 mins |
| 119 Mb | 5 | F<sub>1</sub> (ABxCC) | 10 | 40 | 0.2% | 189,329 | 10,000 | 7 | 5 | 99.8% | 0.985 | 6 hrs 5 mins |
| 119 Mb | 5 | F<sub>1</sub> (ABxCC) | 10 | 30 | 0.2% | 179,218 | 10,000 | 7 | 5 | 99.8% | 0.985 | 6 hrs 20 mins |
| 119 Mb | 5 | F<sub>1</sub> (ABxCC) | 10 | 20 | 0.2% | 127,628 | 10,000 | 7 | 5 | 99.8% | 0.985 | 6 hrs 0 mins |
| 119 Mb | 5 | F<sub>1</sub> (ABxCC) | 10 | 10 | 0.2% | 112,655 | 10,000 | 7 | 5 | 99.8% | 0.986 | 5 hrs 40 mins |
| iii) Altering parental heterozygosity. |  |  |  |  |  |  |  |  |  |  |  |  |
| 119 Mb | 5 | F<sub>1</sub> (ABxCC) | 10 | 50 | 0.01% | 9,996 | 9,996 | 7 | 5 | 99.8% | 0.985 | 6 hrs 49 mins |
| 119 Mb | 5 | F<sub>1</sub> (ABxCC) | 10 | 50 | 0.1% | 97,446 | 10,000 | 7 | 5 | 99.8% | 0.984 | 7 hrs 3 mins |
| 119 Mb | 5 | F<sub>1</sub> (ABxCC) | 10 | 50 | 0.2% | 186,523 | 10,000 | 7 | 5 | 99.8% | 0.986 | 6 hrs 13 mins |
| 119 Mb | 5 | F<sub>1</sub> (ABxCC) | 10 | 50 | 0.5% | 401,020 | 10,000 | 7 | 5 | 99.8% | 0.983 | 7 hrs 6 mins |
| 119 Mb | 5 | F<sub>1</sub> (ABxCC) | 10 | 50 | 1% | 736,293 | 10,000 | 7 | 5 | 99.6% | 0.984 | 6 hrs 40 mins |
| 119 Mb | 5 | F<sub>1</sub> (ABxCC) | 10 | 50 | 2% | 1,148,814 | 10,000 | 7 | 5 | 99.2% | 0.984 | 7 hrs 12 mins |
| iv) Altering genome size and chromosome number. |  |  |  |  |  |  |  |  |  |  |  |  |
| 119 Mb | 5 | F<sub>1</sub> (ABxCC) | 10 | 50 | 0.2% | 186,523 | 10,000 | 7 | 5 | 99.8% | 0.986 | 6 hrs 13 mins |
| 500 Mb | 19 | F<sub>1</sub> (ABxCC) | 10 | 50 | 0.2% | 490,397 | 50,000 | 7 | 5 | 99.2% | 0.984 | 12 hrs 23 mins |
| 900 Mb | 9 | F<sub>1</sub> (ABxCC) | 10 | 50 | 0.2% | 801,712 | 25,000 | 7 | 5 | 99.3% | 0.982 | 22 hrs 0 mins |
| v) Altering cross type |  |  |  |  |  |  |  |  |  |  |  |  |
| 119 Mb | 5 | F<sub>1</sub> (ABxCC) | 10 | 50 | 0.2% | 186,523 | 10,000 | 7 | 5 | 99.8% | 0.986 | 6 hrs 13 mins |
| 119 Mb | 5 | F<sub>2</sub> (AAxCC) | 10 | 50 | 0% | 94,428<br>104,106 | 10,000<br>10,000 | 7<br>7 | 5<br>5 | 100%<br>>99.9% | 0.978<br>0.977 | 7 hrs 49 mins |

AFLAP analysis of an F_2_ population of *Arabidopsis thaliana* using WGS data AFLAP was then validated on real sequencing data using a F_2_ population generated from A. *thaliana* Col x Ler that had previously been sequenced to low coverage (1x to 8x; Additional File 2: Table S1) and analyzed genetically (9-11). Based on the distributions of 31-mers from both parents, the boundaries for classification as a single copy 31-mer were defined as 32 to 105x for *A. thaliana* Col and 41 to 177x for *A. thaliana* Ler (Figure 2a), totaling 109,276,920 and 114,129,149 single copy 31-mers, respectively. The unique number of single copy 31-mers were 21,704,129 for Col and 27,891,980 for Ler.

**Figure 2.**
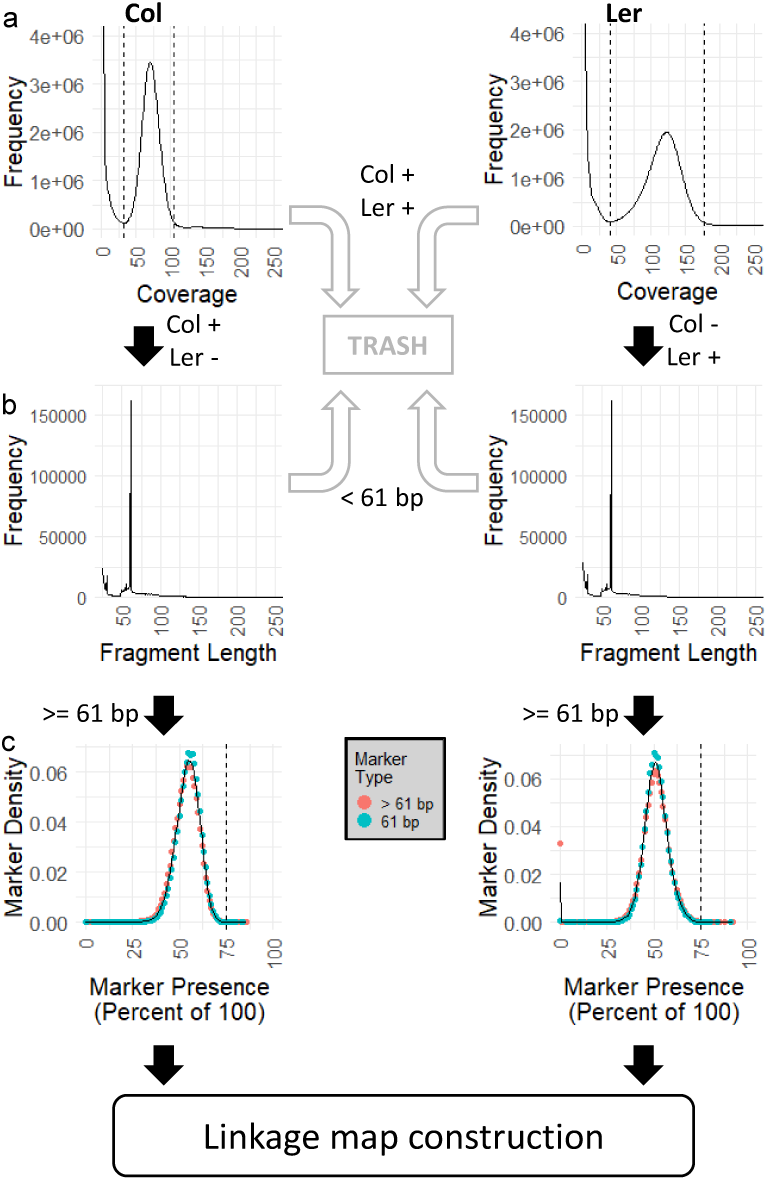
Intermediate data produced by AFLAP when analyzing an *A. thaliana* F_2_ population. Analysis of parental accession Colombia (Col) is plotted on the left and Landsberg (Ler) on the right. a) JELLYFISH histograms were plotted to determine the lower and upper boundaries of single copy 31-mers (indicated by dotted lines). 31-mers found in both parents were discarded (grey arrows) from further analysis. 31-mers unique to either accession were assembled for each parent (black arrows). b) The distribution of the assembled fragments from 31-mers showing that most of these fragments are 61 bp (SNPs). Assembled fragments <61 bp were discarded (grey arrows). A representative 31-mer marker was extracted from each fragment ≥61 bp, verified against the parents, and used for downstream genotyping (black arrows). b) Segregation of 31-mer markers in the F_2_ population. Markers extracted from fragments >61 bp segregate at the same frequency as markers derived from fragments exactly =61 bp, although >3% of Ler derived markers from fragments >61 bp were not observed in the F_2_ population. The modal segregation of Colombia markers was 55%, for Landsberg it was 51%. The black vertical line indicated the expected marker presence of 75% in the F2 (Aa x Aa).

Assembly of unique, single-copy Col 31-mers resulted in 499,936 fragments ranging from 25 to 6,324 bp. Of these, 162,206 fragments were 61 bp and had a SNP in the middle at their 31^st^ base; 156,106 fragments were larger than 61 bp, representing more complex variants; and 181,624 fragments were smaller than 61 bp and contained repetitive or low complexity sequences that were difficult to assemble and were therefore not used in downstream analyses because they represented potentially unreliable variants (Figure 2b). 31-mers were extracted from the 318,312 fragments ≥61 bp, 285,492 (89.9%) of which were confirmed to be within the single copy limits of Col and absent from Ler. On average, these 31-mer markers were scored as present in 55% of F_2_ progeny (Figure 2c).

Assembly of unique, single-copy Ler 31-mers resulted in 519,493 fragments ranging from 25 bp to 26,666 bp. Of these, 161,955 fragments equaled 61 bp (SNPs), 161,174 fragments were larger than 61 bp (complex variants), and 196,364 fragments were <61 bp (Figure 2b). 31-mers were extracted from the 323,129 fragments ≥61 bp, 321,026 (99.3%) of which were confirmed to be within the single copy limits of Ler and absent from Col. On average, these 31-mer markers were scored as present in 51% of F_2_ progeny (Figure 2c). Additionally, 5,329 markers, nearly all of which were derived from markers >61 bp, were not detected in any F_2_ plants (Figure 2c). Only 13 of these markers could be aligned to the reference genome assembly, consistent with these markers being derived from contaminant reads in the SRA dataset for Ler.

Of the 285,492 markers derived from Col, 50.9% were ordered into 315 genetic bins across five linkage groups using a logarithm of the odds (LOD) score ≥ 7. The total map length was 379 cM and linkage groups ranged from 64 cM to 86 cM (Figure 3a). Of the genetically placed markers, 144,395 markers (99.4%) were aligned to the *A. thaliana* reference assembly. Over 99.99% of the markers were concordant with the physical map (Figure 3b). Of the 321,026 markers derived from Ler, 68.0% were ordered into 366 genetic bins across six linkage groups (LOD ≥7). The number of markers assigned to each genetic bin ranged from 1 to 5,857. The total map length was 421 cM and linkage groups ranged from 53 cM to 98 cM (Figure 3c). Of the genetically placed markers, 138,434 (63.4%) were placed unambiguously on the *A. thaliana* reference assembly, with 98.8% concordance between the genetic map and the physical assembly. Two linkage groups mapped to Chromosome 3, each covering a chromosome arm (Figure 3d).

**Figure 3.**
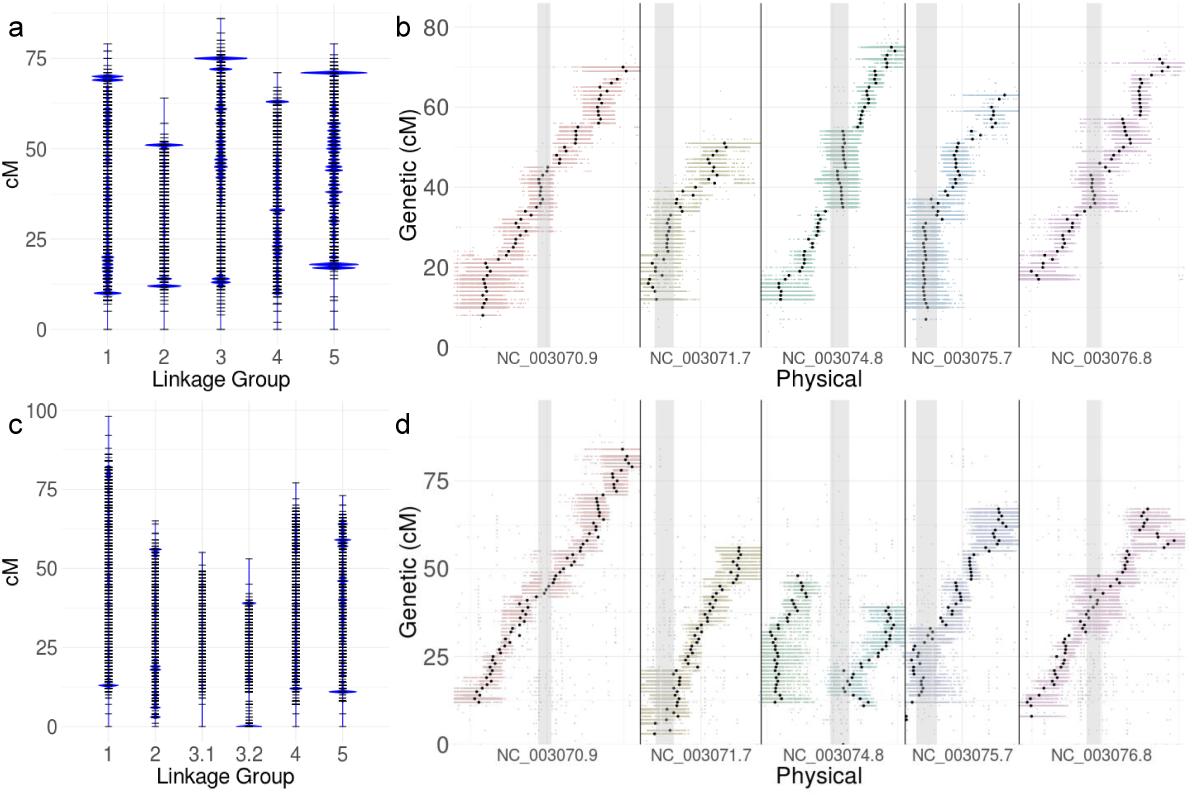
Genetic map of *A. thaliana* generated by AFLAP. a) 145,330 markers unique to *A. thaliana* accession Col ordered into five linkage groups. Each horizontal black bar indicates a genetic bin with >1 marker. The sizes of the blue marks indicate the number of markers in each bin relative to every other genetic bin. Genetic bins with the highest marker densities were near the ends of linkage groups. b) Alignment of the genetic map constructed from markers unique to Col against the genome assembly of Col-0. All 31-mers assigned to each genetic bin were aligned to the assembly and plotted. Points are colored by linkage group assignment resulting in a bar depicting the noise for placement of each genetic bin. The average position of each genetic bin is plotted in black, demonstrating that the linkage groups were nearly colinear with the physical assembly. Grey columns indicate centromeric positions. Over 99.99% of the markers were placed on the correct chromosomes and were concordant with the assembly. c) 115,810 markers unique to *A. thaliana* accession Ler ordered into six linkage groups. Sizes of blue marks indicate the number of markers per genetic bin. d) Alignment of the genetic map constructed from markers unique to Ler displayed as for Col (B). Arms of Chromosome 3 were generated as independent linkage groups. 98.8% of the markers were placed on the correct chromosomes and were nearly concordant with the assembly. In both b and d, the inexact placement of markers into genetic bins demonstrated by the range of the colored bars is probably due to missing data because of the low coverage of the sequencing data.

AFLAP analysis of a RIL population of *Lactuca* spp. using GBS data AFLAP was run on 235 F6 RILs generated by crossing Armenian999 (*L. serriola*) and PI251246 (*L. sativa*), which had previously been sequenced using GBS with 249.8 Mb generated for each line (12). It was not possible to obtain the single copy boundaries from GBS data as had been calculated for *A. thaliana*. Instead, limits were set to ≥20x and ≤45x read depths so markers would not be derived from under- or over-represented sequences. After filtering Armenian999 against PI251246 31-mers, 496,010 unique 31-mers remained. These were assembled into 3,681 markers; 631 of which were 61 bp and 3,050 were <61 bp. The reverse, filtering PI251246 31-mers against Armenian999 identified 650,616 unique 31-mers belonging to PI251246 that were assembled into 5,264 markers; 915 of which were 61 bp and 4,349 were <61 bp. The assembly step also produced many potential markers <61 bp for both parents, which were not used in the subsequent analysis. Markers ≥61 bp segregated approximately 1:1, as expected, indicating that the markers derived from GBS reads by AFLAP were robust (Figure 4a,b). Linkage analysis with Armenian999-derived markers produced a nine-linkage group, 1,730 cM genetic map, containing 3,241 markers (88% of the total identified) placed in 1,656 genetic bins (Figure 4c). Linkage analysis with PI251246-derived markers also produced a nine-linkage group genetic map of 1,705 cM, containing 4,947 markers (94% of the total identified) placed in 2,037 genetic bins (Figure 4d). Both parental maps were aligned to the *L. sativa* cultivar (cv.) Salinas genome assembly (13) to determine if they were colinear. Unique alignments were found for 747 Armenian999-derived markers, across 602 genetic bins in the map (Figure 4e). For PI251246-derived markers, unique alignments for 2,235 markers placed in 1,290 genetic bins were identified (Figure 4f). Both maps were colinear with the genome assembly. As the RILs were largely homozygous, a compound map could be calculated by combining the genotype calls for PI251246-derived and Armenian999-derived markers. The compound map was 1711 cM across nine linkage groups, containing 8,191 markers in 2,497 genetic bins. The compound map was also colinear with the genome assembly, with unique alignments found for 2,984 markers across 1,556 genetic bins (Additional File 3: Figure S1). Therefore, AFLAP can be used to analyze RIL populations and can effectively genotype individuals using GBS data.

**Figure 4.**
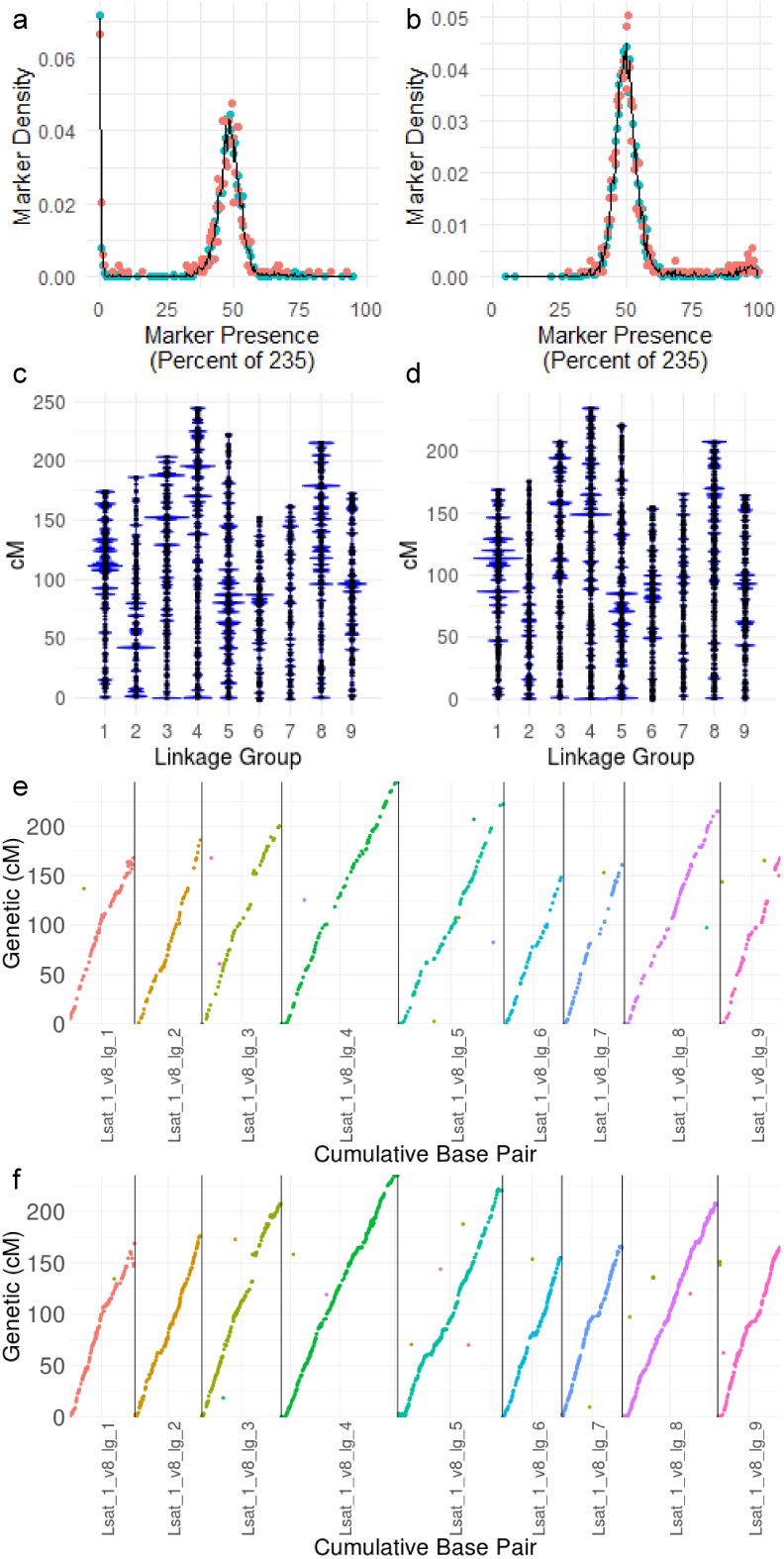
Genetic maps of *Lactuca* spp. produced by AFLAP. Parental and 235 F_6_ recombinant inbred lines of an interspecific *L. serriola* x *L. sativa* cross were analyzed using genotyping-by-sequencing. The segregation of markers derived from a) Armenian999 (*L. serriola*) and b) PI251246 (*L. sativa*) peaked at approximately 0.5, as expected for an F_6_ RIL population. c) The female genetic map was 1,730 cM and contained 3,241 Armenian999-derived markers in 1,656 genetic bins across nine linkage groups. d) The male genetic map was 1,705 cM and contained 4,947 PI251246-derived markers placed in 2,037 genetic bins across nine linkage groups. Unique alignments of e) Armenian999-derived markers and f) PI251246-derived markers along the nine chromosomes of *L. sativa* genome demonstrated collinearity with the genome. Because these RILs were predominantly homozygous, a compound map could also be calculated from a combined genotype table (Additional File 3: Figure S1).

AFLAP analysis of an F_1_ population of *Bremia lactucae* using WGS data AFLAP was then used to genetically analyze the obligately biotrophic oomycete *Bremia lactucae*, for which there was only a partial genetic map and incomplete genome assembly.

Eighty-three F_1_ progeny isolates that had been generated by crossing *B. lactucae* isolate SF5 to either isolate C82P24 or isolate C98O622b (5, 6) were whole genome sequenced to greater than 5x coverage. Based on the distributions of 31-mers from isolate SF5 of *B. lactucae* (Figure 5a), the boundaries for classification as a single-copy, heterozygous 31-mer were 63x to 123x, identifying 27,691,779 potentially useful 31-mers. When compared to the 31-mer compositions of the other parental isolates, C82P24 and C98O622b, 591,159 informative 31-mers were found to be unique to SF5.

**Figure 5.**
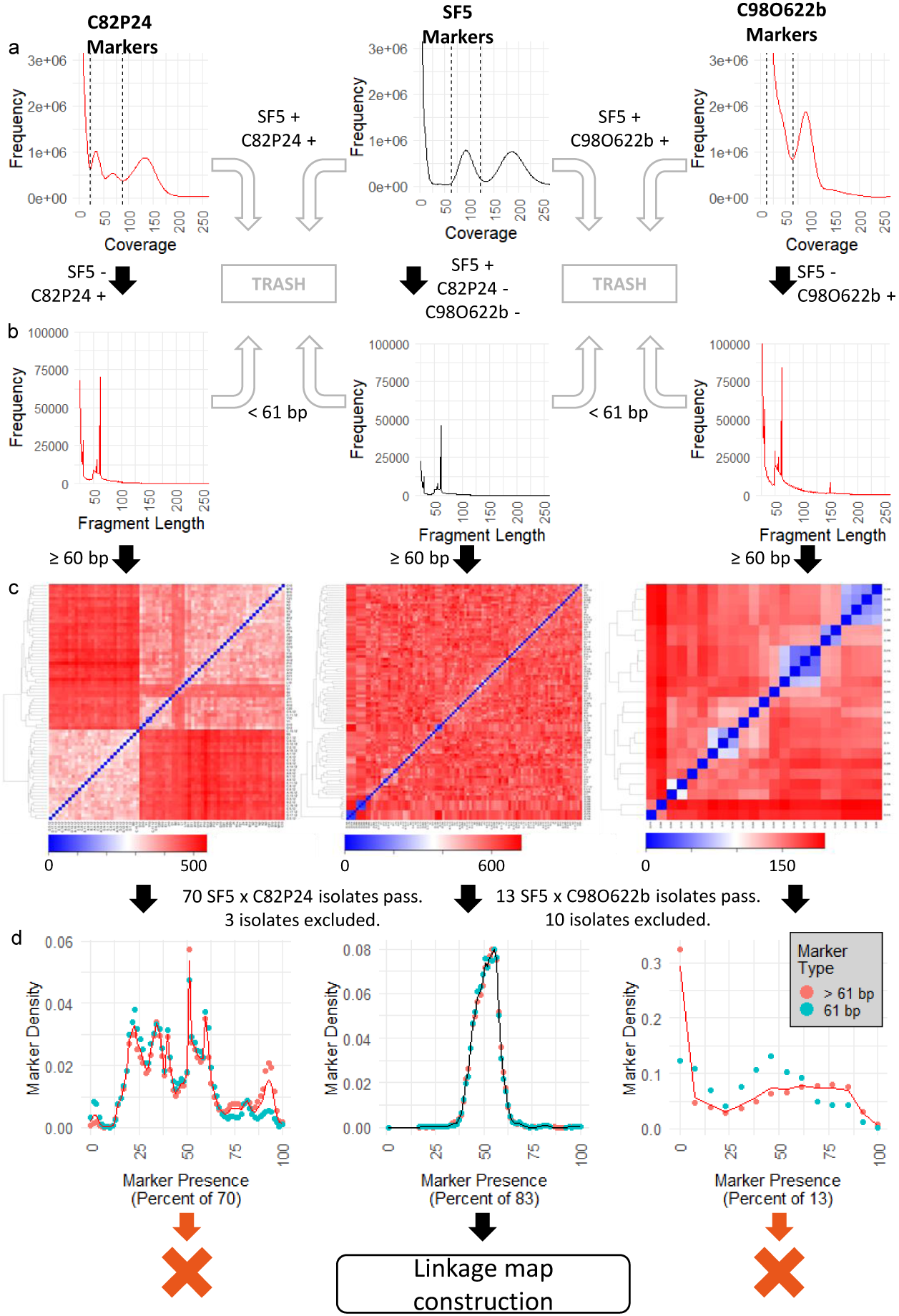
Intermediate data produced by AFLAP when analyzing two F_1_ populations of *B. lactucae*. Two populations generated by crossing SF5 (center) to C82P24 (left) or C98622b (right). Black arrows indicate the path of 31-mers derived from each parental isolate. Ultimately only those derived from SF5 were used for linkage analysis. a) JELLYFISH histograms were generated to determine the boundaries of single copy, heterozygous 31-mers, (dashed lines). 31-mers common to both parents of each cross were discarded (grey arrows). Unique 31-mers to each parental isolate were assembled (vertical arrows). b) The modal assembled fragment size was 61 bp (SNPs), except for C98O622b. Fragments <61 bp were discarded (grey arrows). One representative 31-mer marker was extracted from each fragment ≥61 bp, verified against the parents, and used for downstream genotyping (vertical arrows). c) Kinship heatmaps for parent specific, pseudo-test cross markers from each parent. Left: 73 progeny isolates from SF5 x C82P24 tested with C82P24 derived markers ≥61 bp. Center: 96 progeny isolates from both crosses tested with SF5 derived markers ≥61 bp. Right: 23 progeny isolates from SF5 x C98O622b tested with C98O622b derived markers equal to 61 bp. One isolate was selected when clusters of isolates (colored blue) were formed using markers of both parents. The additional pattern observed for C82P24 (left) is due to heterokaryosis (see 6). d) Segregation of 31-mer pseudo-test cross markers in the F_1_. For C82P24 (left) and SF5 (center), both marker types ≥61 bp segregate at the same frequency. The discordance observed for markers derived from C98O622b may be due to contamination (see a). Distortion is observed for C82P24 because this isolate is heterokaryotic (see c and 6). SF5 is homokaryotic so all gametes are derived from a single nucleus. Therefore, only SF5 derived markers were suitable for linkage map construction.

Assembly of heterozygous 31-mers unique to SF5 resulted in fragments ranging from 25 bp to 2,686 bp. Of these, 45,849 fragments equaled 61 bp representing a SNP in the middle at their 31^st^ base; 59,712 fragments were larger than 61 bp, representative of more complex variants; 132,323 fragments were smaller than 61 bp, representing potentially unreliable variants including repetitive or low complexity sequences that were difficult to assemble (Figure 5b). The set of 31-mer markers unique to SF5 was extracted from the 105,561 fragments equal to or greater than 61 bp, 103,246 (97.8 %) of which were confirmed to be within the heterozygous boundaries of isolate SF5 and absent from the two other parental isolates.

The same process was repeated for the two heterokaryotic parental isolates, C82P24 and C98O622b, to analyze kinship. For C82P24, boundaries of 22x to 88x were identified after visual inspection of the 31-mer distribution (Figure 5a), totaling 40,209,579 31-mers. Compared to the SF5 hash, 19,179,719 were unique to C82P24. Assembly of the unique, heterozygous 31-mers resulted in fragments ranging from 25 bp to 20,022 bp. Of these, 69,868 fragments were 61 bp (SNPs), 123,926 fragments were >61 bp (complex variants), and 335,149 fragments were <61 bp (unreliable variants; Figure 5b). Of the 193,794 markers ≥61 bp, 190,753 (98.4 %) were confirmed to be absent in SF5 and heterozygous in C82P24. For C98O622b, manual inspection of the 31-mer distribution curve indicated that it was not possible to differentiate the lower limits of the heterozygous 31-mer peaks from contaminant sequences (Figure 5a) resulting from sequencing xenic cultures of the biotrophic *B. lactucae*. Therefore, 142,669,686 31-mers with a lower limit of 12x and an upper limit of 65x were selected as the heterozygous component.

When compared to the SF5 hash, 117,091,383 31-mers were found to be unique to C98O622b, which assembled into fragments ranging from 25 bp to 59,784 bp. Of these, 84,435 fragments were 61 bp, 512,948 fragments were >61 bp, and 836,933 fragments were <61 bp (Figure 5b). Because of the high count and inability to resolve 31-mers heterozygous to C98O622b from those belonging to contaminant organisms, only the representative markers of C98O622b SNPs (equal to 61 bp) were used for further analysis. Of these 84,435 markers, 72,496 (85.9 %) were absent in SF5 and heterozygous in C98O622b.

Kinship was analyzed by clustering unique and heterozygous markers to identify near-identical progeny isolates that shared highly similar genotypes. From 73 sexual progeny generated by crossing SF5 by C82P24, six were consistently identified as duplicates with low Euclidean distances between them when considering markers derived from both parents (Figure 5c). One of each pair was omitted from downstream analysis (Additional File 2: Table S2). Fifteen of the 23 progeny generated by crossing SF5 by C98O622b clustered into one of five replicate groups, each with high within group similarity, based on both SF5 and C98O622b markers (Figure 5c). Consequently, ten isolates were excluded, and five representative isolates were used for downstream analysis (Additional File 2: Table S2). This resulted in 83 isolates available for further analysis (Additional File 3: Figure S2).

Additional structure is visible in the kinship analysis of heterokaryotic isolates C82P24 and C98O622b. For C82P24, two large sub-populations of progeny, consisting of 28 progeny (bottom left) and 42 progeny isolates (excluding duplicates, top right), can be identified (Figure 5c); this reflects the heterokaryotic nature of C82P24, where two nuclei contribute independent sets of gametes to the progeny (6). The same pattern can be seen in C98O622b (Figure 5c), which is also a heterokaryon, although only two isolates make up the first sub population (bottom left) and 11 isolates the second (excluding replicates, top right). These patterns are not observed in the SF5 markers (Figure 5c) because it is a homokaryon and therefore only contributes one set of gametes. Finally, for C82P24, four isolates in the second heterokaryon group appear to cluster with one another at a greater Euclidean distance from the rest of the progeny (Figure 5c). The reason for this clustering is unclear because clustering of these isolates was not observed with SF5 markers (Figure 5c); therefore, these isolates were retained for downstream analysis.

Presence of markers in progeny isolates was used to filter for segregation distortion. On average, SF5 31-mer markers were scored as present in 55% of the 83 F_1_ progeny. A total of 5,845 markers (5.6 %) present in ≤33 or ≥52 isolates were filtered out due to segregation distortion (Figure 5d). Therefore, 97,401 markers were used for linkage analysis. Subsequent analysis that increased the cut-off values from 0.4–0.6 to 0.2–0.8 did not greatly increase the number of markers nor did it result in more sequence being captured in the linkage map; this reflects the selection of the boundaries as indicated in Figure 5d.

Markers originating from SF5 were ordered into 1,337 genetic bins across 19 linkage groups, placing 98.8% (96,226) of the markers. The total map length was 1,769 cM and linkage groups ranged from 52 cM to 148 cM (Figure 6a). Of these markers, 61.4% (59,087) aligned unambiguously to the previously reported *B. lactucae* assembly, 96.9% (57,247) of which aligned to scaffolds larger than 1 Mb. This accounted for 96.6% of the 115.9 Mb genome assembly (6). Long stretches of genetic markers were colinear with this assembly; however, 19 of the 21 scaffolds larger than 1 Mb contained sequences from different linkage groups and therefore appeared to be chimeric (Figure 6b). Chimeras may have resulted from false joins due to the highly repetitive architecture of the *B. lactucae* genome (6). The chimeric scaffolds were broken using the linkage data. Reorienting and re-scaffolding resulted in 97 Mb organized into 19 scaffolds, each compassing a single linkage group (Figure 6c). The remaining 18.6 Mb compromising 200 scaffolds did not have genetic markers aligned and so remained unplaced. This included six scaffolds over 1 Mb. Repeat-masking revealed that 70% of the genetically placed contig sequence and 73% of the unplaced contig sequence was repetitive. A higher percentage of the unplaced large scaffolds were covered by C82P24 and C98O622b derived markers than SF5 derived markers (Additional File 3: Figure S3). Of the 9,767 annotated genes, 8,345 were placed into the 19 large scaffolds and 1,422 were on genetically unassigned scaffolds; 261 out of a total 280 candidate effector genes were located across all linkage groups, except linkage group 17 (Figure 6c). The dark diagonal obtained when analyzing Hi-C contact frequency demonstrated that the genetically orientated assembly was consistent with the Hi-C data (Figure 6d). Attempts to refine the assembly using Hi-C reads and scaffolding software did not improve the assembly. Synteny of the genetically-oriented assembly of *B. lactucae* with *Phytophthora sojae* showed that the gene order between the two oomycete assemblies was highly conserved (Figure 7). Therefore, scaffolding using the AFLAP genetic map was able to correct errors in the genome assembly of *B. lactucae* and produce linkage-group-scale scaffolds that are coherent with Hi-C data and largely syntenic with a distantly related oomycete.

**Figure 6.**
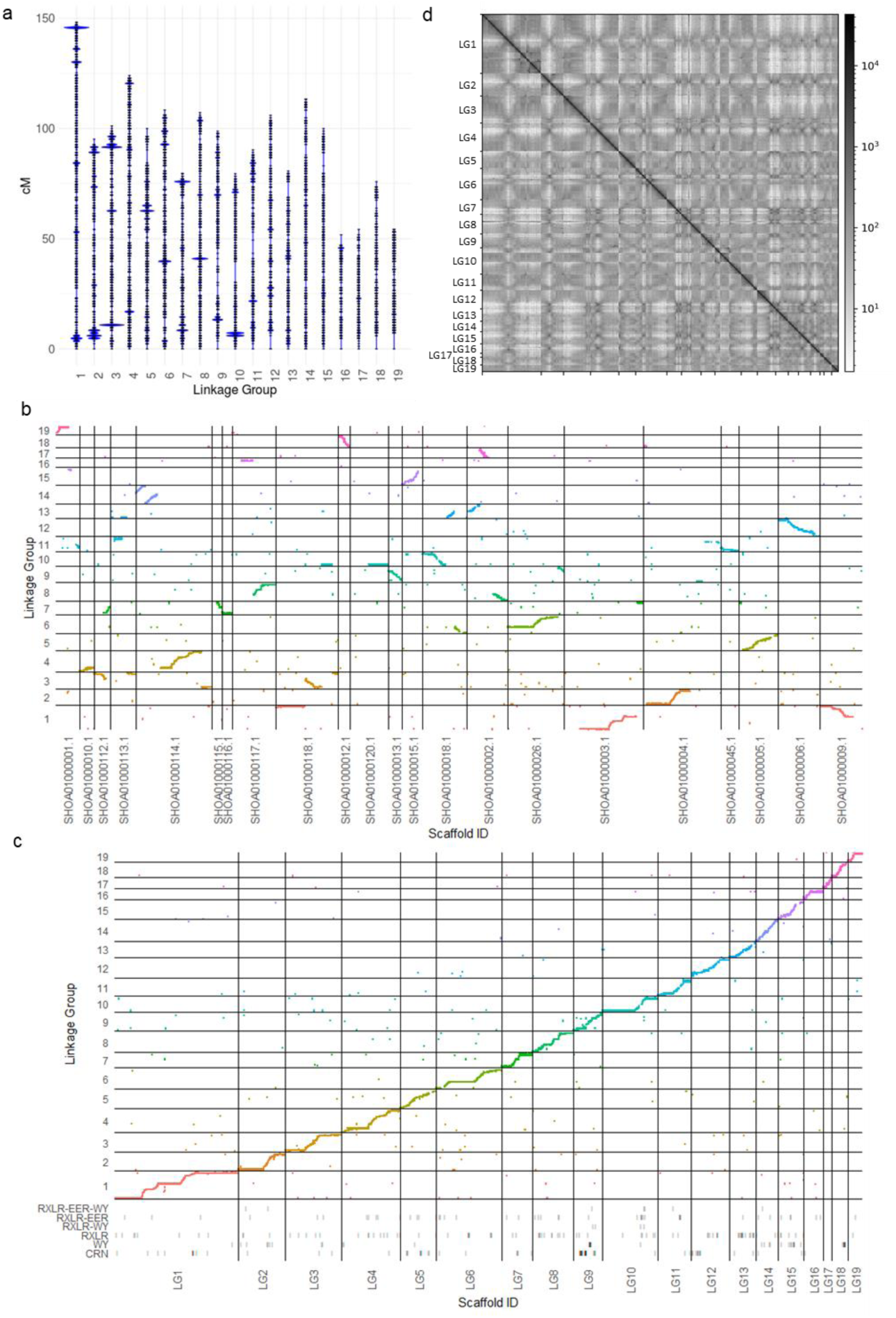
*Bremia lactucae* AFLAP results. a) Markers unique to SF5 ordered in 19 linkage groups. Each horizontal bar indicates a genetic bin with >1 marker. The sizes of the blue marks indicate the relative number of markers in each genetic bin. Genetic bins with large numbers of markers were observed at the end of some, but not all linkage groups. b) Alignment of the genetic map on the 21 scaffolds larger than 1 Mb in the draft reference assembly (6). Multiple scaffolds spanned more than one linkage group, indicative of mis-assembly. Linkage groups included multiple scaffolds providing genetic support for reorienting and rejoining the scaffolds. c) Scatter plot demonstrating collinearity between the revised genome assembly and the genetic map after fragmenting, reorienting, and re-scaffolding of the assembly. Coordinates of annotated genes that encode six categories of candidate effector proteins are plotted on tracks below the scatter plot. d) Contact frequency obtained by aligning paired Hi-C reads to the genetically reoriented assembly. The strong dark diagonal indicates that the assembly is consistent with Hi-C data. Crosses off the diagonal indicate intra-chromosomal contacts, possibly involving centromeres.

**Figure 7.**
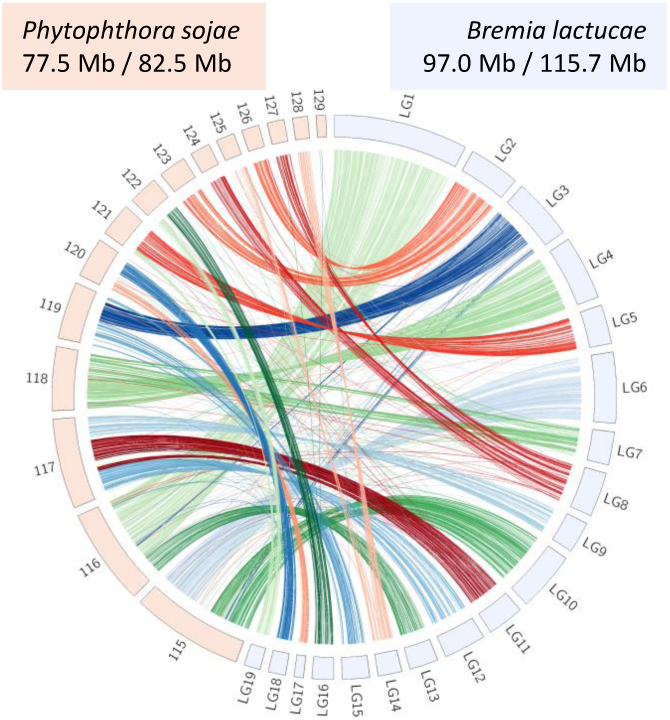
Synteny between *Bremia lactucae* and *Phytophthora sojae*. Single copy orthologs of *B. lactucae* and *P. sojae* were used to link genetically revised scaffolds of *B. lactucae* (light blue) with scaffolds of *P. sojae* larger than 1 Mb (pink). *P. sojae* scaffolds are labelled with the three-digit suffix of their NCBI accession (i.e., 115 is NW_009258115.1). Links are colored based on their assignment on *B. lactucae* scaffolds. Gene order is highly conserved between the two assemblies providing strong support for the quality of both.

## Discussion

We developed the Assembly Free Linkage Analysis Pipeline, AFLAP, to generate ultra-dense genetic maps based on single copy *k*-mers without reference to a genome assembly. This approach to linkage analysis does not require reads to be mapped and variants called against a reference assembly for marker identification. Instead, variants are identified using assembled *k*-mers, of fixed length *k*, unique and single copy to each parent. Assembled fragments equal to *k + k -* 1 are considered equivalent to SNPs, with the variant position present at the center of the fragment. Fragments larger than *k + k - 1* are likely to be complex variants such as insertions and deletions. Therefore, AFLAP uses markers generated from fragments larger or equal to *k + k - 1*. These fragments are reduced to a representative marker, containing the variant, equal in length to *k* so that a) all markers have the same size and b) a constant marker size can be surveyed in the progeny downstream. AFLAP enables the rapid construction of a genotype table for subsequent linkage analysis.

We tested AFLAP using 100 F_2_ individuals of *A. thaliana* sequenced to low coverage. The genetic architecture of *A. thaliana* has been studied in detail; over 2,000 F_2_ individuals, generated by crossing Colombia (Col) x Landsberg (Ler), have been sequenced to low coverage (9-11). The 100 individuals with the largest number of reads from this population were selected to create a test population of similar size to the total progeny of the two *B. lactucae* experimental populations. The *A. thaliana* markers were expected to segregate in a 1:2:1 ratio; however, this was not the case. The modal percentage of progeny markers detected was 55% for Col markers and 51% for Ler markers (Figure 2c). Missing data can therefore be estimated as between 26% and 32%. Despite this, the size of the genetic map is very similar to that reported previously (14, 15). In addition, the average physical positions of genetic bins were highly concordant with the genome assembly with 99% of the markers assigned to the correct chromosome (Figure 3b, d). The noise in the precise placement of the genetic bins and the low percentage of genetically placed markers (50.9% for Col, 63.4% for Ler derived markers) was likely caused by missing data due to low sequence coverage, as shown with the simulated data (Table 2, Additional File 1). Despite the imperfect input data, AFLAP was able to produce a good genetic map using markers from each parent, concordant with the chromosome-scale genome assembly.

AFLAP was then applied to F_1_ progeny isolates of *B. lactucae* generated by crossing isolate SF5 with either C82P24 or C98O622b, both of which are heterokaryotic; therefore, SF5 was effectively crossed to four different nuclei (6). Progeny isolates were whole genome sequenced to at least 5x coverage to provide reliable identification of unique 31-mers. Heterokaryosis was reflected by half-sib clusters of isolates in the progeny (Figure 5c). In addition to heterokaryosis, clustering of 31-mer markers also demonstrated that some isolates were genetically more similar to other isolates than expected, allowing potential duplicate individuals to be removed. Therefore, 83 isolates were genotyped with SF5-specific markers and used for linkage analysis (Additional File 3: Figure S2). The small population sizes for individual nuclei from the heterokaryotic parents meant that maps of C82P24 and C98O622b could not be constructed.

The genetic map of isolate SF5 of *B. lactucae* produced by AFLAP placed 98.8% of the SF5-specific markers into 19 linkage groups (Figure 6a). The genetic map was highly concordant with large portions of the published genome assembly (Figure 6 b; 6). Discordance between the genetic map and the assembly was used to identify mis-assemblies; linkage data was then used to guide binning, orienting, and scaffolding, resulting in a much-improved genome assembly with 19 linkage-group-scale scaffolds (Figure 6c). The more accurate placement of genetic bins on the assembly and higher percentage of mapped makers when compared to *A. thaliana* (Figure 3b, d) is likely due to the higher coverage in the *B. lactucae* dataset. The size of the genetic map produced for *B. lactucae* is similar to that reported previously (4). Therefore, with adequate sequencing depth, AFLAP was able to rapidly produce a genetic map of a non-model organism with a highly repeated genome (6); the high marker density enabled genetically-guided fragmentation and re-scaffolding of the genome assembly.

Not all scaffolds were placed on the linkage map. The marker sparse regions totaling 18.6 Mb of the current assembly were only marginally more repetitive than the genetically oriented sequence. Pseudo-test cross markers derived from isolates C82P24 and C98O622b did align to the large unplaced scaffold in the *B. lactucae* assembly (Additional File 3: Figure S3); therefore, it is possible that the unplaced regions over 1 Mb are homozygous in isolate SF5. In the current study, not enough progeny isolates were obtained from any of the nuclei of the heterokaryotic isolates C82P24 or C98O622b for genetic analysis. Therefore, additional genetic analysis of other isolates will further refine the *B. lactucae* genome assembly. Genotyping more progeny isolates and generating a consensus map will determine if *B. lactucae* has fewer than 19 chromosomes. Aligning the assemblies of *B. lactucae* and *P. sojae* allowed potential joins to be inferred based on synteny (e.g., Figure 7; scaffold 117 of *P. sojae* suggests that linkage groups 9, 11, and 12 of *B. lactucae* might belong to a single chromosome). Alternatively, enhanced genetic resolution may demonstrate that the genomes of these distant relatives have undergone large-scale structural variation since divergence from their common ancestor. It is possible that applying AFLAP to *P. sojae* could further refine the *P. sojae* assembly, investigating syntenic joins suggested by the new assembly of *B. lactucae* (e.g., Figure 7; linkage group 2 of *B. lactucae* joins scaffolds 123 and 127 of *P. sojae*).

Markers derived from fragments under 61 bp were investigated by rerunning AFLAP including markers derived from fragments equaling 60 bp. Many fragments smaller than *k + k - 1* are probably derived from low complexity, repetitive, or hard to assemble sequences and are therefore uninformative. Some fragments equal to 60 bp will contain instances of single base pair deletions at a locus and therefore will be informative (Additional File 3: Figure S4). Rerunning AFLAP including markers derived from 60 bp fragments only added 4,070 markers to the 96,226 markers used to construct the *B. lactucae* map (Figure 6) and did not alter the marker ordering (Additional File 3: Figure S5). Given that the very large number of markers far exceeded the number of crossovers, using markers derived from smaller fragments was unnecessary to generate accurate genetic maps. Indeed, AFLAP can generate robust genetic maps using only markers derived from 61 bp fragments. Depending on the genetics of the organism under study, there may be advantages to including markers derived from smaller fragments or only using markers derived from 61 bp fragments.

AFLAP has several technical benefits over other strategies for linkage analysis. It is not subject to biases that may be introduced by a reference assembly due to reads from reference alleles mapping more readily to an assembly than reads from alternative alleles (16) or associated SNP calling errors. In addition, AFLAP enables access to all single-copy portions of the genomes, some of which may not be present in the reference assembly. This may be particularly important when the parents of the mapping population are distantly related to the reference genotype. AFLAP makes it possible to genotype multi-nucleotide polymorphisms and indels in addition to SNPs; such variants are often overlooked in conventional mapping approaches.

Therefore, AFLAP removes bias in marker calling and increases access to variants and genetic markers. GBS/RADseq can be used to obtain high coverage, reduced representation sequencing data, and using ustacks, linkage analysis may be performed without an assembly (17); however, library preparation for GBS/RADseq may introduce allele bias caused by restriction site distribution (18), a lack of robust genotype calls, and much lower marker density. The utility of AFLAP was demonstrated on a lettuce RIL population genotyped using GBS (12). AFLAP generated nine linkage groups for each parent colinear with the 2.4 Gb genome assembly (Figure 4) (13). A nine-linkage group, 1,711 cM nine-linkage group compound map (Additional File 3: Figure S1) was concordant with a 1,883 cM genetic map previously obtained via a conventional read alignment and variant calling workflow (12). Therefore, AFLAP can efficiently generate accurate genotype tables for linkage analysis from GBS data. AFLAP also allows facile addition of data from new progeny individuals from the same or different populations that have a common parent to increase the genetic resolution of the map. Because each isolate is genotyped independently, adding new isolates generated from the same parents is equivalent to appending additional columns to the genotype table. Adding data from a new cross, but sharing one parent, can be achieved by filtering the 31-mer marker set against the new parent and removing markers from the genotype table that are no longer unique to the common parent. When analyzing the interspecific *Lactuca* spp. RILs (Additional File 3: Figure S1) the compound map was generated by concatenating the genotype table containing *L. serriola* derived markers to the end of the genotype table containing *L. sativa* derived markers (i.e., genotypes did not require recalculation). Therefore, AFLAP can incorporate new data easily, enabling rapid maturation of genetic maps.

AFLAP enables the construction accurate genotype tables resulting in high-quality genetic maps for any organism using a segregating population sequenced to adequate depth. Analyses using simulated and real data demonstrated that the sequence depth obtained on progeny affects the accuracy of marker placement in the genetic map. Even low coverage sequencing (3x) is adequate to assign a marker to a correct linkage group with approximate placement. The accuracy of genetic placement of markers increased as progeny sequencing depth increased (Table 2). The desired sequencing depth will therefore vary depending on the aims of the project. For validation of a chromosome scale assembly, low coverage may be adequate. For genetic orientation of a fragmented assembly, at least 5x coverage in the progeny is required. In simulated data, more markers were required to accurately place markers in genomes containing more chromosomes. AFLAP may be applicable to many datasets already generated or being generated. Also, WGS data generated for AFLAP can be easily repurposed for use in numerous other projects. Workflows, such as AFLAP, that use unbiased WGS as input will become increasingly desirable as the costs of library generation and sequencing continue to decrease. This may be critical to validating genome assemblies of non-model species generated in projects such as the Earth BioGenome Project (19).

## Conclusions

AFLAP is a novel *k*-mer based approach to linkage analysis able to produce a high-density genetic map, without a prerequisite reference genome assembly. In addition, AFLAP can analyze complex variants and genomic regions that may not be accessible to variant calling approaches that use a reference genome assembly. AFLAP was benchmarked using multiple simulations, varying the sequencing depth of progeny and parents, parental heterozygosity, genome size, and chromosome number and contrasted to a conventional read alignment, variant calling workflow. AFLAP was validated using low coverage *A. thaliana* F_2_ WGS data and was able to construct linkage groups that were coherent to the reference assembly when aligned; however, the use of low coverage data introduced significant noise. The utility of AFLAP when using GBS data was demonstrated by analyzing a RIL population of a *Lactuca* interspecific cross. AFLAP was then deployed to analyze 83 F_1_ isolates of the non-model oomycete *B. lactucae* that had been whole genome sequenced to ≥5x. The genetic maps produced were unambiguously aligned to the reference assembly and resulted in significant improvements of the assembly. AFLAP can therefore be used to generate saturated genetic maps and to improve draft genome assemblies of non-model organisms provided a mapping population with adequate sequencing coverage is available.

## Materials and Methods

### Whole genome sequencing

Whole genome sequencing of *A. thaliana* accessions Colombia (Col) and Landsberg (Ler) were downloaded from NCBI Short Read Archive (SRA) accessions SRR5882797 and SRR3166543 (20), respectively. Low coverage WGS reads of F_2_ individuals generated from Col x Ler were obtained from previous studies (9-11); 100 individuals with the largest gzipped files were selected for analysis (Additional File 2: Table S1). Reads for parents and RILs of a previously analyzed *L. serriola* Armenian999 x *L. sativa* PI251246 interspecific-cross were downloaded from NCBI BioProjects PRJNA642889, PRJNA510128, and PRJNA478460 (12).

Two *B. lactucae* crosses were analyzed with parental isolate SF5 in common. For both crosses, WGS of *B. lactucae* parental isolates SF5, C82P24, and C98O622b have been described previously (6); NCBI BioProject PRJNA387613). Thirty-seven previously reported F_1_ progeny (6); NCBI BioProject PRJNA387454) and an additional 36 F_1_ progeny from the same cross, made earlier (5), were added. Extracted DNA that had previously been used for RFLP linkage analysis (5) was used to construct ∼350 bp libraries using the Lucigen (Middleton, WI, USA) NxSeq HT dual indexing kit, per the manufacturer’s instruction and 150 bp paired end reads were generated on an Illumina NovaSeq 6000 lane. For the second cross, oospores were obtained by co-inoculating isolates SF5 and C98O622b onto cv. Cobham Green. Oospores matured for several weeks in decaying plant tissue prior to maceration. Isolates were generated by growing cv. Cobham Green in a dilute oospore suspension, which was titrated via serial dilution so that on average a single seedling would be infected per culture box. DNA was extracted by vortexing sporangia for two minutes in a microcentrifuge tube with approximately 200 μL of Rainex-treated beads and 0.5 mL of 2× extraction buffer (100 mM Tris-HCl pH 8.0, 1.4 M NaCl, 20 mM EDTA, 2% [wt/vol] cetyltrimethylammonium bromide, and B-mercaptoethanol at 20 μL/mL), then transferred to a fresh 2 mL tube. Material was treated with RNase (20 μL/mL; 65°C for 30 min). An equal volume of 1:1 phenol/chloroform was added, mixed, and centrifuged at maximum speed (8,000 rpm; 15 min). The aqueous phase was retained and further washed twice with equal volumes of 24:1 chloroform/isoamyl alcohol, obtaining the aqueous phase each time by centrifuging at maximum speed for 15 min. The resulting aqueous phase was mixed with 0.7 volumes of isopropanol and DNA was precipitated at −20°C for one hour. DNA was pelleted by centrifuging at maximum speed for 30 min, washing with 70% ethanol, drying, and suspending in 10 mM Tris-HCl. Quantity and quality of DNA was determined by spectrometry, as well as estimated by TAE gel electrophoresis. Single index libraries of 23 F_1_ isolates were generated by sonicating DNA to ∼ 220 bp (Covaris, Woburn, MA, USA), cleaning and concentrating (1 part DNA : 1.2 parts AMPure, Beckman Coulter, Pasadena, CA USA), end repairing (End Repair Module #E6050L, New England Biolabs, Ipswich, MA, USA), cleaning (1 part DNA : 1.2 parts AMPure), A-base ligating (Klenow, Enzymatics/Qiagen, Hilden, Germany), and adapter ligating (T4 DNA Ligase #L603-HC-L, Enzymatics/Qiagen). Final cleanup and size selection was performed using 1-part DNA: 0.8 parts AMPure. Paired end, 150 bp reads were generated by sequencing the libraries with an Illumina HiSeq 4000. Reads new to this study have been deposited under BioProjects PRJNA387454 and PRJNA634525.

### Assembly Free Linkage Analysis Pipeline (AFLAP)

Figure 1 provides an overview of the pipeline. For *A. thaliana, B. lactucae*, and *Lactuca* spp., 31-mer hashes were produced independently for the read sets of each parent and each progeny individual using JELLYFISH (21) sub-command count, parameters -*m31* -C -s *10G*. The parental (F_0_) hashes were then inspected to identify single copy 31-mers using the JELLYFISH sub-command histo to produce histograms of parental hashes, which were manually inspected to determine lower and upper bounds for filtering. For *A. thaliana*, where both parental accessions are highly homozygous and the population analyzed was an F_2_, all single copy 31-mers were retained. For *Lactuca* spp., where both parental lines were sequenced by reduced representation GBS, no single copy 31-mers could be recovered, so user-specified limits of ≥20 to ≤45x were supplied to avoid sampling markers from under- or over-represented sequence. For *B. lactucae*, where both parental isolates are highly heterozygous and the population was an F_1_, only the heterozygous 31-mers from either parent were retained. FASTA files were obtained for each parent using the JELLYFISH sub-command dump, parameters *-L [LowerLimit] -U [UpperLimit]*. For *A. thaliana* and *B. lactucae*, single-copy 31-mers were then queried against the opposite parental hash using JELLYFISH sub-command query and filtered for zero counts. The resulting 31-mers were single copy and homozygous (*A. thaliana*) or heterozygous (*B. lactucae*) in one parent and absent in the alternate parent. For *Lactuca* spp., retained 31-mers were also queried against the opposite parental hash using JELLYFISH sub-command query and filtered for zero counts, so the resulting 31-mers were unique to either parent, although may not be single copy.

To reduce redundancy, 31-mers for each parent were assembled using ABYSS v2.2.2 (22), with the parameters *-k25 -c0 -e0* (23). Assembled fragments equal to or greater than 61 bp were extracted and a single representative 31-mer equal to coordinates 10 to 41 was selected for each fragment, though any 31-mer would have sufficed. Fragments equal to 61 bp were considered single nucleotide variants and therefore were used as markers for the subsequent genetic analysis. Fragments larger than 61 bp were considered complex variants, including insertions and deletions, were also used. Fragments smaller than 61 bp were likely to contain repetitive or low complexity sequences that were not easily assembled and therefore not considered suitable for use as markers. The representative 31-mers were verified against the parental hashes to ensure they were a) within the boundaries set for single copy markers and b) absent in the second parent. This set of markers was then scored against every progeny hash to obtain progeny genotypes. For the low coverage WGS (1x to 8x) *A. thaliana* and GBS *Lactuca* inter-cross lines, the presence of the marker was established by a single observation. For the high coverage (>5x) *B. lactucae*, two or more observations were required for the marker to be scored. The scores were then collated into a genotype table, where 0 = marker not observed and 1 = marker observed.

To filter for siblings of *B. lactucae* with high identity, individuals were clustered by genotype. The Euclidean distance between progeny was calculated for the marker scores using the *dist* function of R (24) package *proxy* (25) These were clustered with the *hclust* function, from which a dendrogram was calculated (*as*.*dendrogram* function) and plotted with the *heatmap*.*2* function from the package gplots (26). Progeny with high identity to one another were reduced to a single representative isolate.

The genotype table was then converted to be compatible with Lep-MAP3 (8). For *A. thaliana* (F_2_ population) and *Lactuca* spp. (RIL population) the F_0_ are coded as grandparents, markers unique to each parent are coded AA in the accession that it was identified in and CC in the alternate parent, for which it was not identified. Inferred, genetically homogeneous F_1_ parents were inserted with every marker coded AC. For F_2_ progeny for which the genotype table was constructed, marker presence was coded AA and marker absence was coded CC. Genotype tables were constructed for markers sourced independently from either parent as well as a combined table from both. For *B. lactucae* (F_1_ population), the identification of a marker was coded AC and marker absence was coded CC in both parents and progeny.

Lep-MAP3 subcommand SeperateChromosomes2 was run on the genotype table to assign markers to linkage groups using the following parameters: *lodLimit=7* for *A. thaliana, lodLimit=20* for *L. sativa*, and *lodLimit=3* for *B. lactucae*. Lep-MAP3 subcommand OrderMarkers2 was subsequently used to order markers within each linkage group using a Morgan mapping function with 20 iterations. The native output file of LepMap3 OrderMarkers2 did not retain the original marker names; instead, these are derived from the genotype table using Linux join. A shell script is provided in the GitHub repository and will generate the output described here. Linkage groups produced were visualized using R (24) packages dplyr (27), ggplot2 (28), and ungeviz (29).

To validate the genetic maps produced by AFLAP, the linkage groups were aligned against corresponding reference assemblies (*A. thaliana*; GCF_000001735.4, *B. lactucae*; GCA_004359215.1, *L. sativa* GCA_002870075.2) by mapping 31-mers to the assembly with bwa aln (30), converting to a sorted BAM file with SAMtools v1.9.1 subcommand sort (31), and filtering for uniquely mapped 31-mers with a maximum one edit distance. Genetic coordinates were visualized across the physical assembly as scatter plots in R using ggplot2 (28), colored by linkage group. For *A. thaliana*, the average physical position of each genetic bin was calculated and plotted due to the low coverage of the F_2_ individuals.

For the simulations, parents and progeny were simulated for multiple organisms representing different genome sizes. The genome assemblies of *A. thaliana* (119 Mb, five chromosomes, NCBI: GCF_000001735.4), *Vitis riparia* (500 Mb, 19 chromosomes, NCBI: GCA_004353265.1), and *Atriplex hortensis* (900 Mb, nine chromosomes, CoGe Genome ID: 56906) were used to simulate F_1_ crosses where markers only segregate from one haplotype (ABxCC). For all assemblies, SNPs and indels were introduced to produce a synthetic parent (AB) containing 0.2 % heterozygosity using mutate.sh (32). The other parent was 100% homozygous and represented by the reference assembly. The impact of varying heterozygosity was tested at levels approximate to 0.01%, 0.1%, 0.2%, 0.5%, 1%, and 2% using the smallest simulated genome. Progeny were generated by randomly assigning an inherited haplotype and either one or two cross-overs along each chromosome (https://github.com/kfletcher88/CrossSimulator). Parental sequencing depth was simulated to 50x and progeny sequencing depth to 10x whole genome coverage using randomreads.sh (32). Simulations were also run varying the parental sequencing depth to 10x, 20x, 30x, and 40x and the progeny sequencing depth to 3x, 5x, 7x, and 20x. F_2_ crosses were simulated using the smallest genome assembly simulating both parents to be 100% homozygous, varying from one another by 0.1%, and synthesizing reads for the parents at 50x and the progeny at 10x. All AFLAP simulations were run using a sub sampled marker set using the shell script AFLAP.sh available at https://github.com/kfletcher88/AFLAP/archive/v0.02.tar.gz.

To compare AFLAP with a contemporary SNP based pipeline, the simulated paired-end reads for the smallest genome assembly were mapped back to the reference assembly using BWA mem (v0.7.17) (33) and SNPs were called using FreeBayes (v1.3.1) (34). The VCF file was recoded using VCFtools (v0.1.16) and converted into a LepMap3 compatible file using the template R-code available at https://github.com/rkbhan/GeneticsTools.git. LepMap3 was run using the entire marker set and a down-sampled subset for comparison with AFLAP. All simulated data was run in series using 12 threads writing to a scratch drive enabling time comparisons. The same crossover coordinates were used for these simulations, so that the map length calculated by AFLAP and the conventional pipelines were comparable. A minimum LOD score of seven was used for AFLAP runs with both the full marker set and the down-sampled marker set. For the conventional run, a minimum LOD score of seven was used for the down-sampled marker set. For the full marker set, a minimum LOD score of 20 was required to resolve the five linkage groups; lower minimum LOD scores resulted in some linkage groups being erroneously joined to one another. Correlation of each synthetic genetic map with the original genome assembly, from which the synthetic data was derived, was inferred by plotting the genetic coordinates by physical coordinates and calculating the Kendall Rank Coefficient (35). All computations were done on the UC Davis Genome Center computing cluster.UC Davis Genome Center computing cluster.

### Curation of the reference genome assembly of *B. lactucae*

Linkage data was used to identify and break chimeric scaffolds outside of gene boundaries followed by linkage-guided reorientation and rejoining of genetically congruent scaffolds. To identify chimeric scaffolds, continuous runs of genetically placed markers were quantified along each scaffold. Spurious results, where a run of less than 10 markers were identified on a scaffold, were disregarded. Scaffolds were broken on the final marker of a run of 10 or more markers, or beyond the boundaries of any genes that were identified to contain the marker. A gene was retained in the linkage group when there was at least one marker providing evidence for the segregation of that gene with the linkage group. The average genetic position of each scaffold was determined based on the genetic position of markers mapped to it. Scaffolds upon which recombination could be detected were oriented based on the average physical position of the genetic bins mapped to each scaffold. Oriented scaffolds were joined with a string of 100 Ns. Unresolved marker sparse or void regions were excised from the scaffold and retained as unlinked scaffolds within the assembly file. Genetic markers were remapped onto the genetically oriented assembly for validation. Hi-C reads previously generated from the same isolate (6); BioProject PRJNA387613) were aligned to the genetically oriented assembly and contact frequencies visualized with Hi-C explorer v2.2 (36) to validate the assembly. Repeats were masked with RepeatMasker v4.0.9 (37) and a previously defined repeat library (6). Genic annotations were lifted over to the new assembly using Liftoff v1.3.0 (38). Annotations that did not transfer or were found to be erroneously transferred were manually corrected. Single copy orthologs previously identified (6) between *B. lactucae* and *Phytophthora sojae* (GCF_000149755.1) were used to plot synteny with circos v0.69-8 (39).

## Additional Files

Additional File 1. Plots of simulated genetic maps aligned back to their respective assemblies, in support of Table 1 and Table 2. (.pdf)

Additional File 2. Table S1. A pedigree file which could be used to run AFLAP, includes accession numbers for *A. thaliana* WGS reads used. Table S2. High identity *B. lactucae* progeny isolates identified in Figure 5 C. (.xls)

Additional File 3. Figure S1. Compound and parental maps of *Lactuca* spp. generated with AFLAP. Figure S2. Corrected *B. lactucae* kinship clustering heatmap. Figure S3. Percent of bases covered by *B. lactucae* pseudo-test cross markers derived from each parent. Figure S4. Associating variants with different fragment sizes. Figure S5. Addition of markers derived from 60 bp fragments to the SF5 genetic map. (.pdf)

## Declarations

### Ethics approval and consent to participate

N/A

### Consent for publication

N/A

Availability of data and materials: All *Bremia* data reported in this paper are available from NCBI BioProjects PRJNA387613, PRJNA387454, PRJNA387192, and PRJNA634525 (this study and 6). Previously published *A. thaliana* reads are available at NCBI short read archive SRR3166543 and SRR5882797 (20) and EBI ArrayExpress E-MTAB-4657, E-MTAB-5476, and E-MTAB-8165 (9-11). Previously published *Lactuca* reads are available in NCBI BioProjects PRJNA642889, PRJNA510128, and PRJNA478460 (12). Scripts to run AFLAP are available at https://github.com/kfletcher88/AFLAP.

## Authors’ contributions

KF conceptualized the project, performed the bioinformatic analysis, and drafted the manuscript. LZ and JG produced isolates of *B. lactucae* and prepared libraries for sequencing. RH conducted the GBS analysis and contributed code to the project. KC prepared libraries of additional isolates. RM supervised the project, contributed to data analysis and all drafts. All authors contributed to writing and editing the paper and have approved the final submission.

## Acknowledgements

We thank B. Rowan (UC Davis) for discussion of results produced by AFLAP on the *A. thaliana* data, W. J. Palmer (UC Davis) for discussions on linkage analysis software to integrate into the pipeline, H Xu (UC Davis) for managing raw data submissions and E. Georgian (UC Davis) for editorial services. The sequencing was carried out by the DNA Technologies and Expression Analysis Cores at the UC Davis Genome Center, supported by NIH Shared Instrumentation Grant 1S10OD010786-01. The bioinformatic analysis was carried out using the UC Davis LSSC0 High Performance Computing cluster maintained by the UC Davis Bioinformatics Core.

