## Additional File 1 for "AFLAP: Assembly-Free Linkage Analysis Pipeline using *k*-mers from whole genome sequencing data"

Contents:

2. Benchmarking AFLAP against a conventional linkage analysis pipeline, in support of Table 1.

3. Simulations exploring the impact of varying progeny sequencing coverage when running AFLAP, in support of Table 2i.

4. Simulations exploring the impact of varying parental sequencing coverage when running AFLAP, in support of Table 2ii.

5. Simulations exploring the impact of varying parental heterozygosity when running AFLAP, in support of Table 2iii.

6. Simulations exploring the impact of varying genome size and chromosome number when running AFLAP, in support of Table 2iv.

7. Simulations exploring the impact of varying cross structure when running AFLAP, in support of Table 2v.

Benchmarking AFLAP against a conventional linkage analysis pipeline, in support of Table 1.

**AFLAP:** 119 Mb, 5 chromosome genome assembly. 10x progeny coverage. 50x parental coverage. 0.2% heterozygosity. **186,523 markers**. Minimum LOD=7.
Time = 59 hours 32 minutes. τ = 0.986

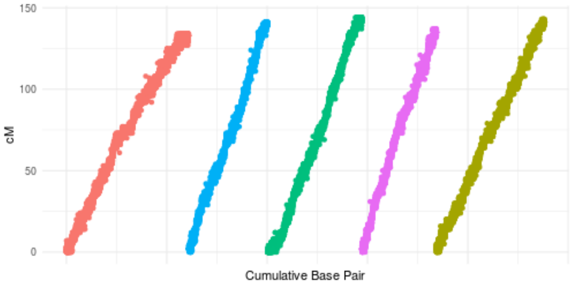


**AFLAP**: 119 Mb, 5 chromosome genome assembly. 10x progeny coverage. 50x parental coverage. 0.2% heterozygosity. **10,000 markers**. Minimum LOD=7.
Time = 6 hours 13 minutes. τ = 0.986

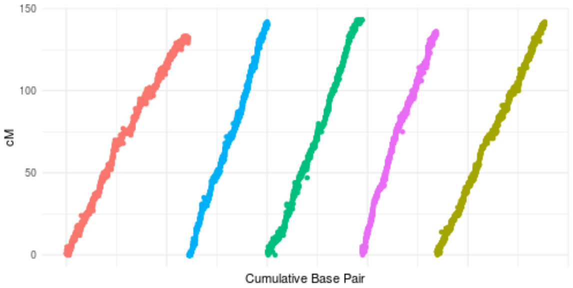


**Read alignment + variant calling pipeline**: 119 Mb, 5 chromosome genome assembly. 10x progeny coverage. 50x parental coverage. 0.2% heterozygosity. **225,797 markers**. Minimum LOD=20.
Time = 173 hours 34 minutes. τ = 0.992

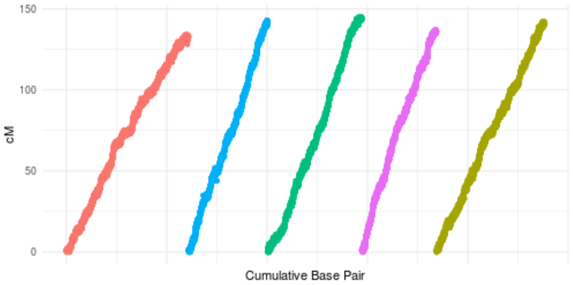


**Read alignment + variant calling pipeline**: 119 Mb, 5 chromosome genome assembly. 10x progeny coverage. 50x parental coverage. 0.2% heterozygosity. **10,000 markers**. Minimum LOD=7.
Time = 107 hours 5 minutes. τ = 0.992

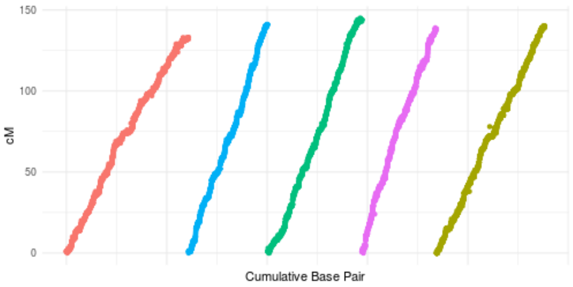

Simulations exploring the impact of varying progeny sequencing coverage when running AFLAP, in support of Table 2i.

AFLAP: 119 Mb, 5 chromosome genome assembly. **20x progeny coverage**. 50x parental coverage. 0.2% heterozygosity. 10,000 markers. Minimum LOD=7.
Time = 8 hours 44 minutes. τ = 0.993

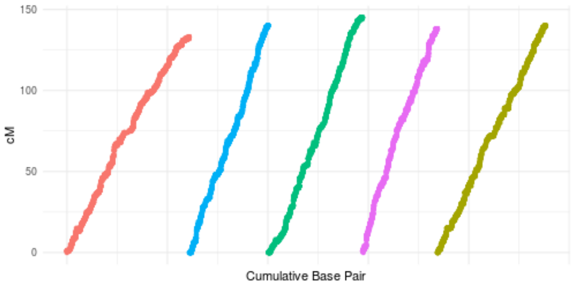


AFLAP: 119 Mb, 5 chromosome genome assembly. **10x progeny coverage**. 50x parental coverage. 0.2% heterozygosity. 10,000 markers. Minimum LOD=7.
Time = 6 hours 13 minutes. τ = 0.986

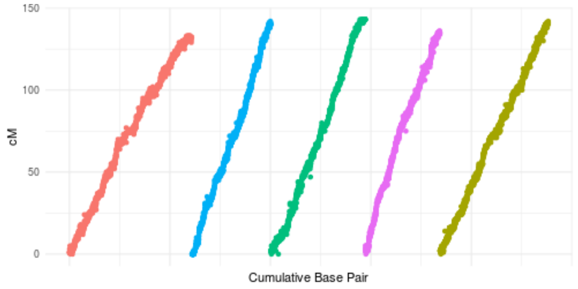


AFLAP: 119 Mb, 5 chromosome genome assembly. **7x progeny coverage**. 50x parental coverage. 0.2% heterozygosity. 10,000 markers. Minimum LOD=7.
Time = 5 hours 18 minutes. τ = 0.988

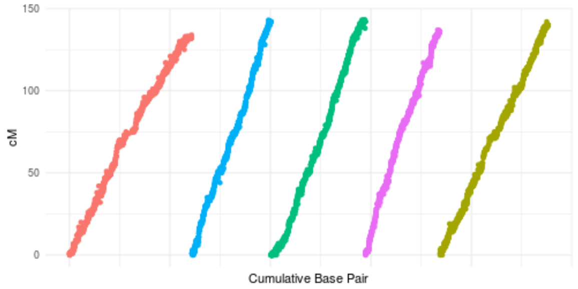


AFLAP: 119 Mb, 5 chromosome genome assembly. **5x progeny coverage**. 50x parental coverage. 0.2% heterozygosity. 10,000 markers. Minimum LOD=7.
Time = 4 hours 22 minutes. τ = 0.983

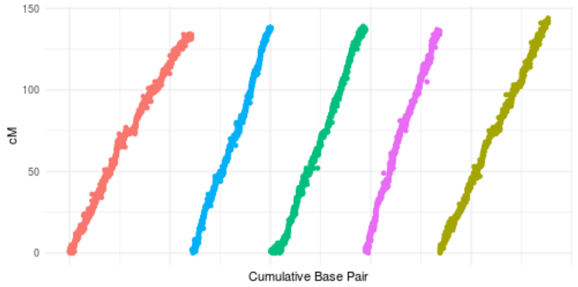


AFLAP: 119 Mb, 5 chromosome genome assembly. 3**x progeny coverage**. 50x parental coverage. 0.2% heterozygosity. 10,000 markers. Minimum LOD=3.
Time = 4 hours 17 minutes. τ = 0.826

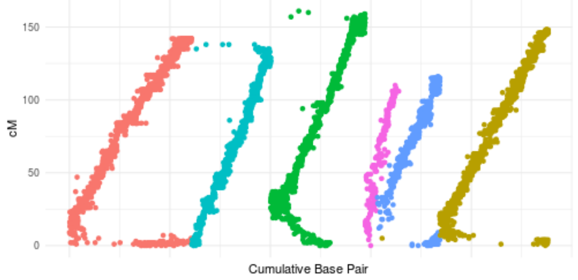

Simulations exploring the impact of varying parental sequencing coverage when running AFLAP, in support of Table 2ii.

AFLAP: 119 Mb, 5 chromosome genome assembly. 10x progeny coverage. **50x parental coverage**. 0.2% heterozygosity. 10,000 markers. Minimum LOD=7.
Time = 6 hours 13 minutes. τ = 0.986

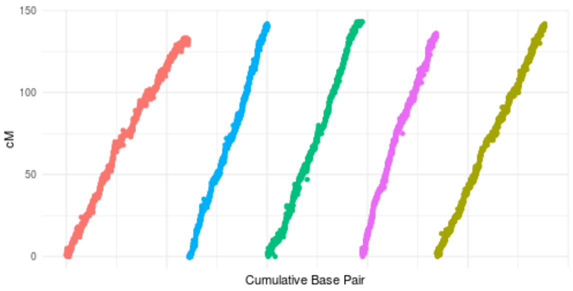


AFLAP: 119 Mb, 5 chromosome genome assembly. 10x progeny coverage. **40x parental coverage**. 0.2% heterozygosity. 10,000 markers. Minimum LOD=7.
Time = 6 hours 5 minutes. τ = 0.985

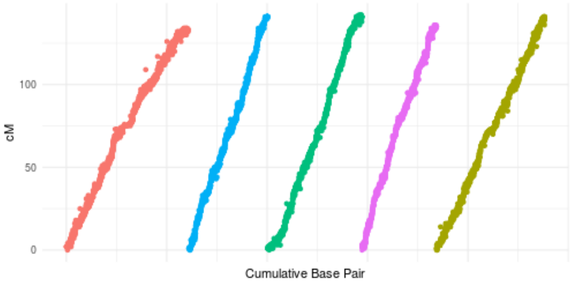


AFLAP: 119 Mb, 5 chromosome genome assembly. 10x progeny coverage. **30x parental coverage**. 0.2% heterozygosity. 10,000 markers. Minimum LOD=7.
Time = 6 hours 20 minutes. τ = 0.985

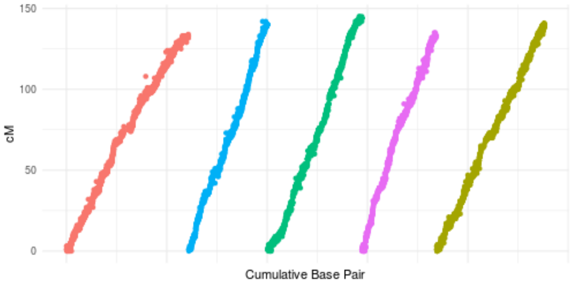


AFLAP: 119 Mb, 5 chromosome genome assembly. 10x progeny coverage. **20x parental coverage**. 0.2% heterozygosity. 10,000 markers. Minimum LOD=7.
Time = 6 hours 0 minutes. τ = 0.985

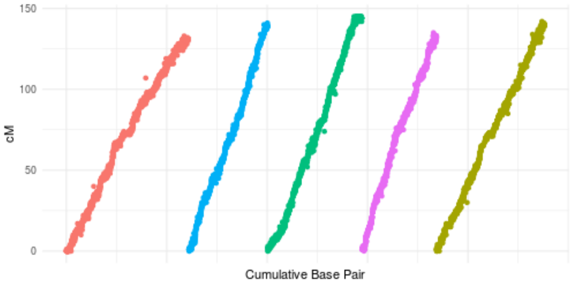


AFLAP: 119 Mb, 5 chromosome genome assembly. 10x progeny coverage. **10x parental coverage**. 0.2% heterozygosity. 10,000 markers. Minimum LOD=7.
Time = 5 hours 40 minutes. τ = 0.986

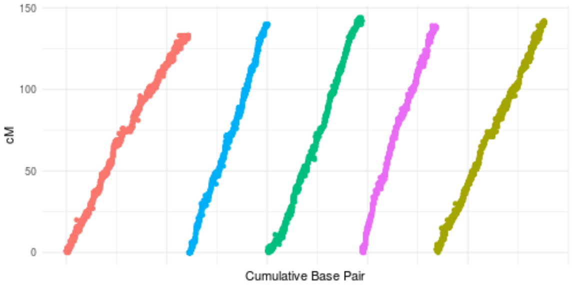

Simulations exploring the impact of varying parental heterozygosity when running AFLAP, in support of Table 2iii.

AFLAP: 119 Mb, 5 chromosome genome assembly. 10x progeny coverage. 50x parental coverage. **0.01% heterozygosity**. 9,996 markers. Minimum LOD=7.
Time = 6 hours 49 minutes. τ = 0.985

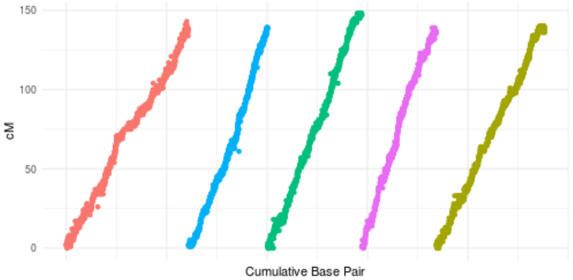


AFLAP: 119 Mb, 5 chromosome genome assembly. 10x progeny coverage. 50x parental coverage. **0.1% heterozygosity**. 10,000 markers. Minimum LOD=7.
Time = 7 hours 4 minutes. τ = 0.984

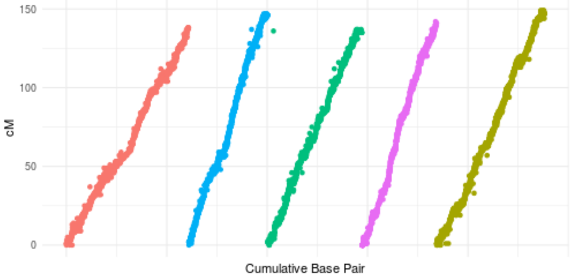


AFLAP: 119 Mb, 5 chromosome genome assembly. 10x progeny coverage. 50x parental coverage. **0.2% heterozygosity**. 10,000 markers. Minimum LOD=7.
Time = 6 hours 13 minutes. τ = 0.986

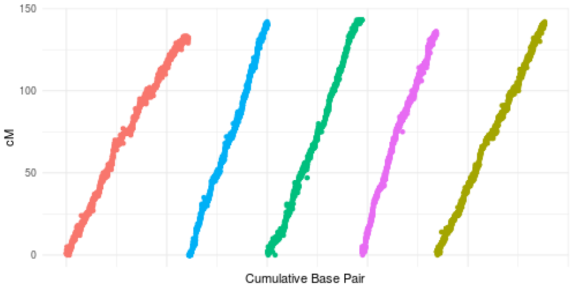


AFLAP: 119 Mb, 5 chromosome genome assembly. 10x progeny coverage. 50x parental coverage. **0.5% heterozygosity**. 10,000 markers. Minimum LOD=7.
Time = 7 hours 6 minutes. τ = 0.983

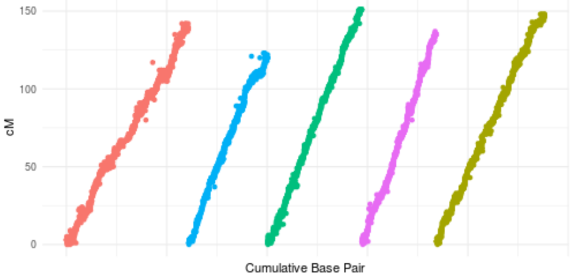


AFLAP: 119 Mb, 5 chromosome genome assembly. 10x progeny coverage. 50x parental coverage. **1% heterozygosity**. 10,000 markers. Minimum LOD=7.
Time = 6 hours 40 minutes. τ = 0.984

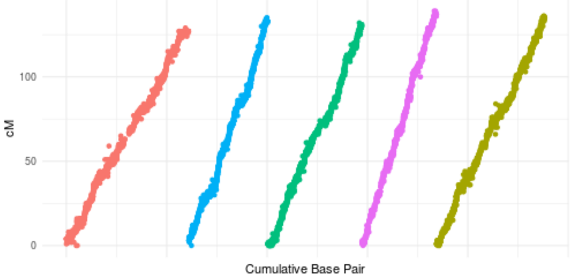


AFLAP: 119 Mb, 5 chromosome genome assembly. 10x progeny coverage. 50x parental coverage. **2% heterozygosity**. 10,000 markers. Minimum LOD=7.
Time = 7 hours 12 minutes. τ = 0.984

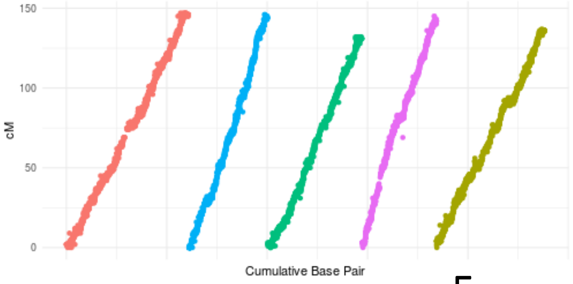

Simulations exploring the impact of varying genome size and chromosome number when running AFLAP, in support of Table 2iv.

AFLAP: **119 Mb, 5 chromosome genome assembly.** 10x progeny coverage. 50x parental coverage. 0.2% heterozygosity. 10,000 markers. Minimum LOD=7.
Time = 6 hours 13 minutes. τ = 0.986

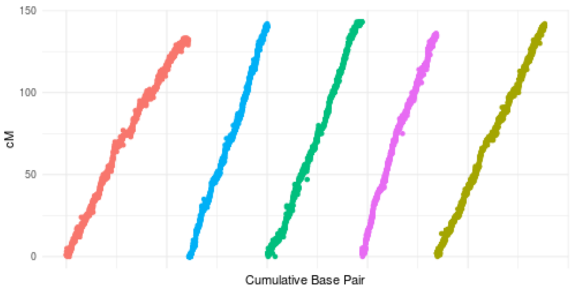


AFLAP: **500 Mb, 19 chromosome genome assembly**. 10x progeny coverage. 50x parental coverage. 0.2% heterozygosity. 50,000 markers. Minimum LOD=7.
Time = 12 hours 23 minutes. τ = 0.984

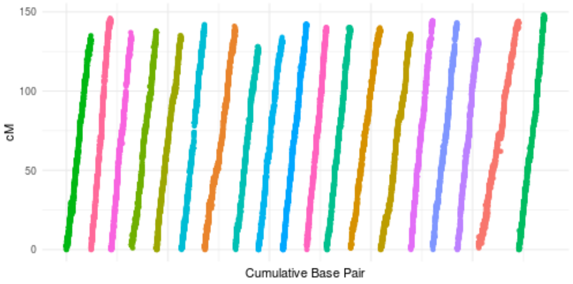


AFLAP: **900 Mb, 9 chromosome genome assembly**. 10x progeny coverage. 50x parental coverage. 0.2% heterozygosity. 25,000 markers. Minimum LOD=7.
Time = 22 hours 0 minutes. τ = 0.982

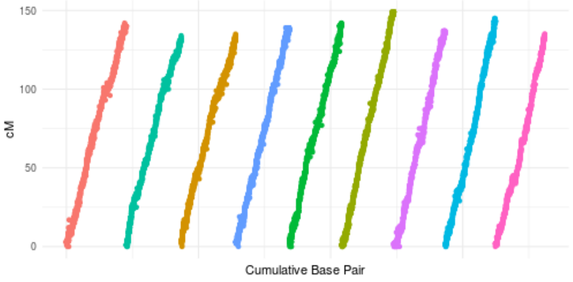

Simulations exploring the impact of varying cross structure when running AFLAP, in support of Table 2v.

AFLAP: 119 Mb, 5 chromosome genome assembly. **$F_1$ cross.** 10x progeny coverage. 50x parental coverage. 0.2% heterozygosity. 10,000 markers. Minimum LOD=7.
Time = 6 hours 13 minutes. τ = 0.986

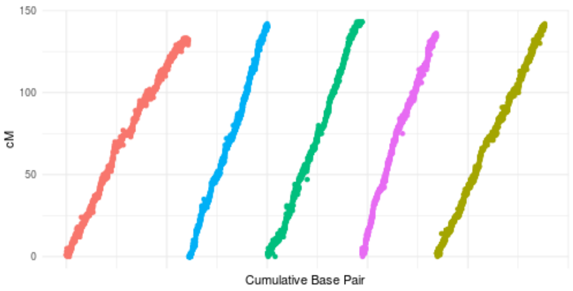


AFLAP: 119 Mb, 5 chromosome genome assembly. **$F_2$ cross.** 10x progeny coverage. 50x parental coverage. 0.2% heterozygosity. 10,000 markers. Minimum LOD=7.
Time = 7 hours 49 minutes.
Parent 1 Map τ = 0.978
Parent 2 Map τ = 0.977

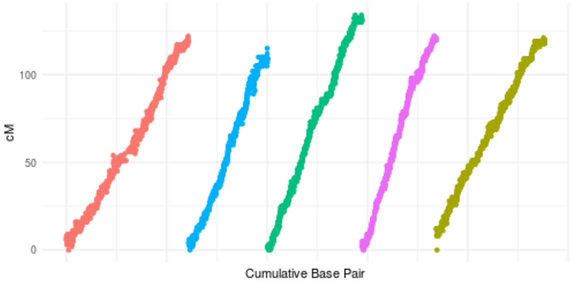


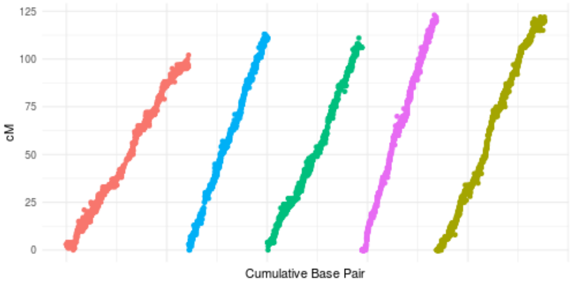
