## Additional File 3: Figures S1 to S5 for "AFLAP: Assembly-Free Linkage Analysis Pipeline using *k*-mers from whole genome sequencing data"

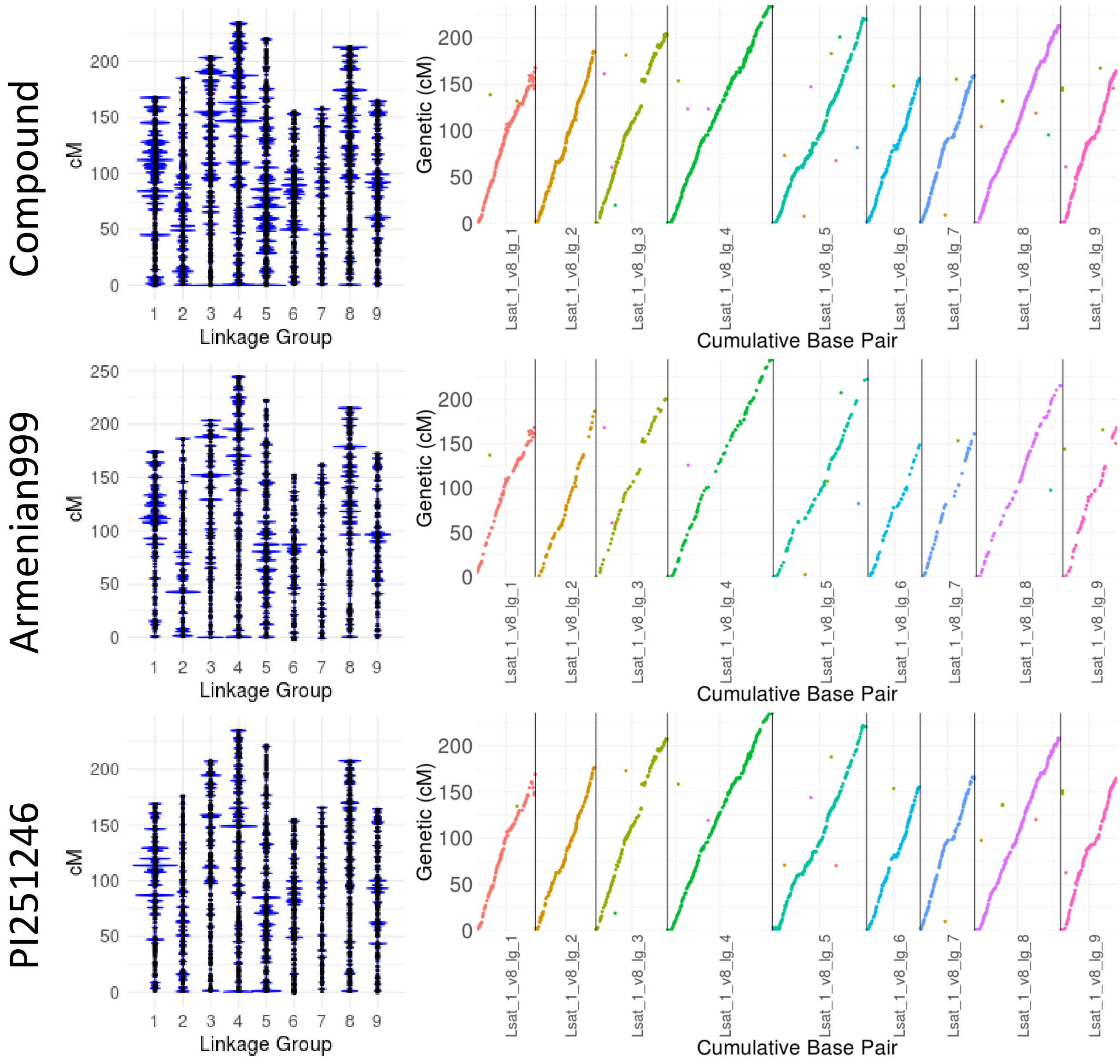


**Figure S1. Comparisons of the compound genetic map with independent *L. serriola* and *L. sativa* genetic maps.** The compound genetic map was 1,711 cM, ordering 8,191 in 2,497 genetic bins along nine linkage groups. Unique alignments against the *L. sativa* assembly were found for 2,984 markers across 1,556 genetic bins and showed collinearity of the genetic map and genome assembly. Armenian999 (*L. serriola*) and PI251246 (*L. sativa*) are repeated from Figure 4 c-f for comparison. The Armenian999 map was 1,705 cM containing 4,947 in 1,656 ordered into nine linkage groups. Unique alignments were found for 747 markers against the *L. sativa* assembly. The PI251246 specific map was 1,730 cM containing 4,947 markers in 2,037 genetic bins ordered into nine linkage groups. Unique alignments were found for 2,235 markers.

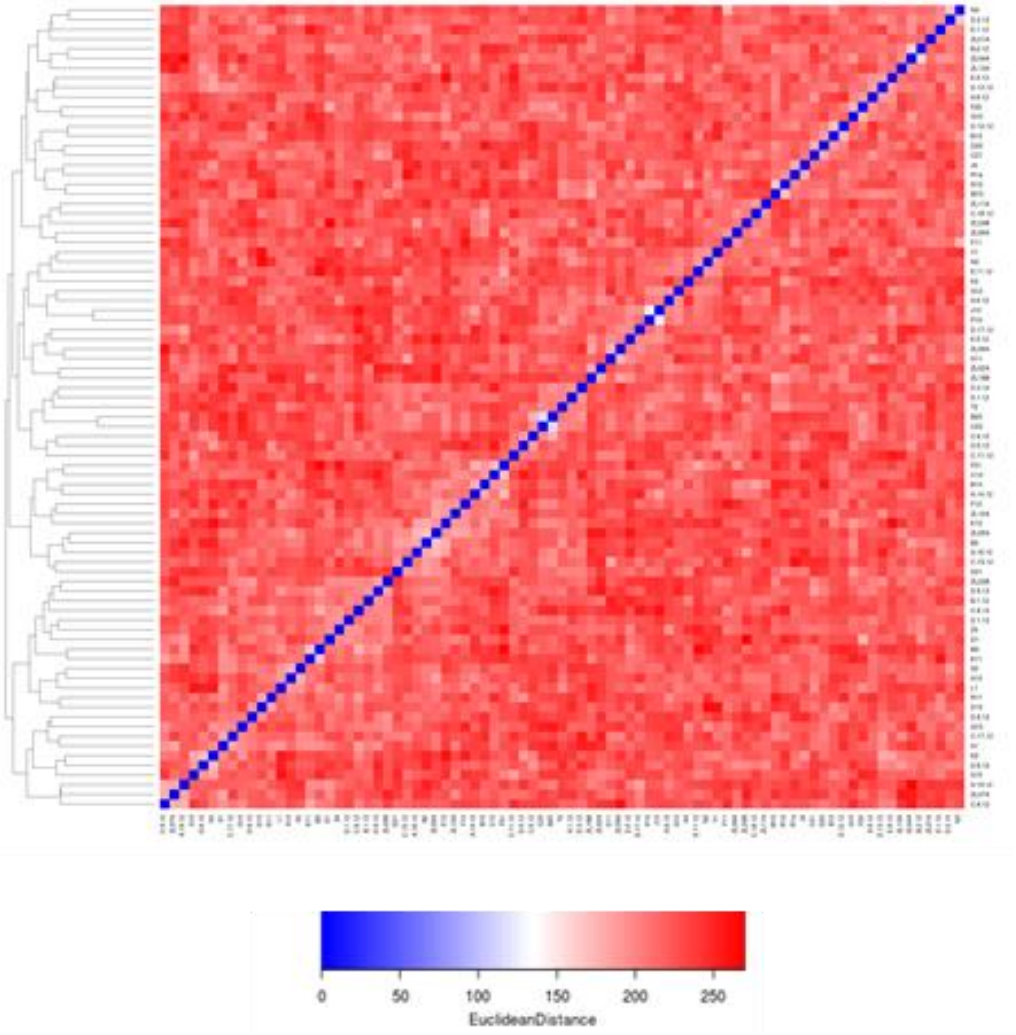


**Figure S2. Corrected *B. lactucae* kinship clustering heatmap.** Heatmap used to infer kinship based on SF5 derived markers ≥61 bp. This figure differs from Figure 4c because high identity clusters of isolates have each been reduced to single representative isolates.

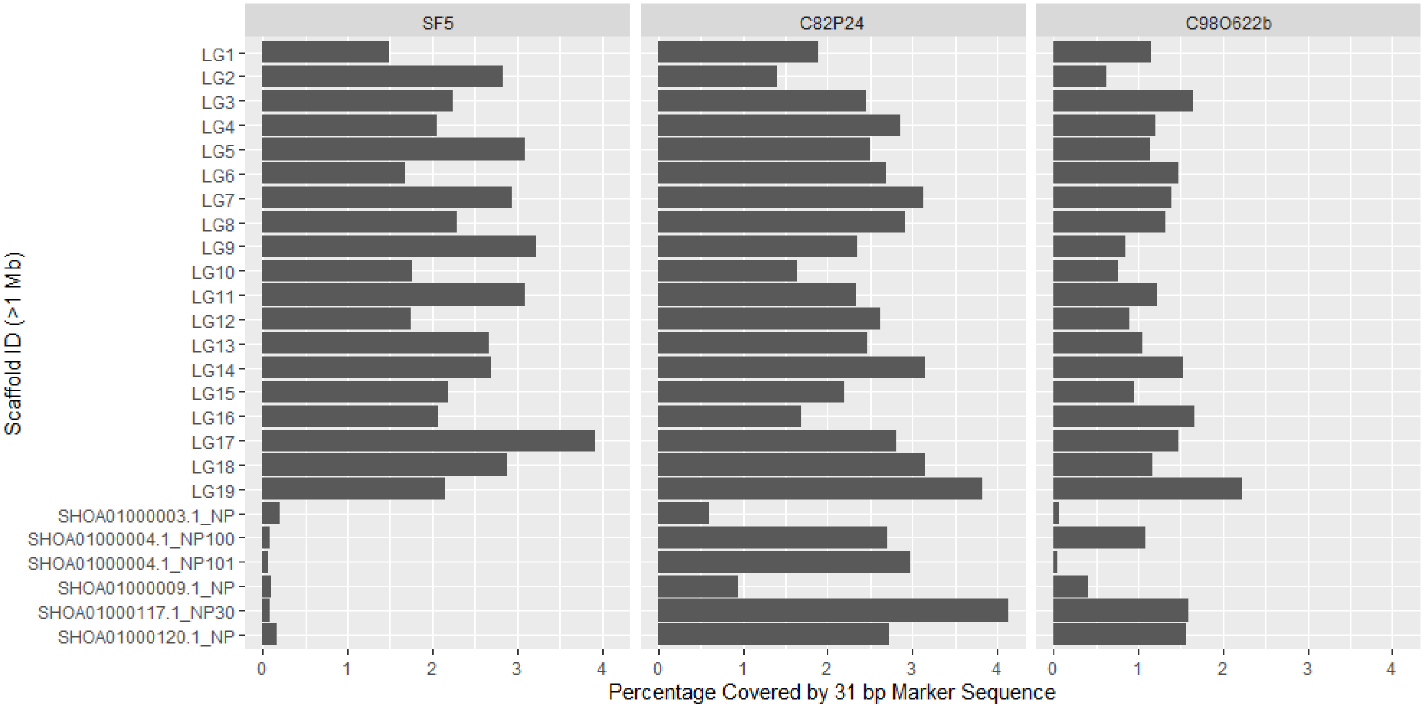


**Figure S3. Percent of bases covered by *B. lactucae* pseudo-test cross markers derived from each parent.** Markers derived from fragments ≥61 bp for SF5 and C82P24 or =61 bp for C98O622b were mapped to the genetically oriented genome assembly. Percentage coverage of scaffolds larger than 1 Mb was plotted. Very few pseudo-test cross markers derived from SF5 aligned to the unplaced scaffolds (prefixed SHOA) compared to genetically oriented scaffolds (prefixed LG) that did. This pattern did not hold for the other parents; some unplaced scaffolds had at least as much coverage as placed scaffolds. It is therefore likely that these scaffolds represent regions of high homozygosity in SF5 but not in other isolates. Therefore, generation of genetic maps of other isolates of *B. lactucae* will further refine the genome assembly.

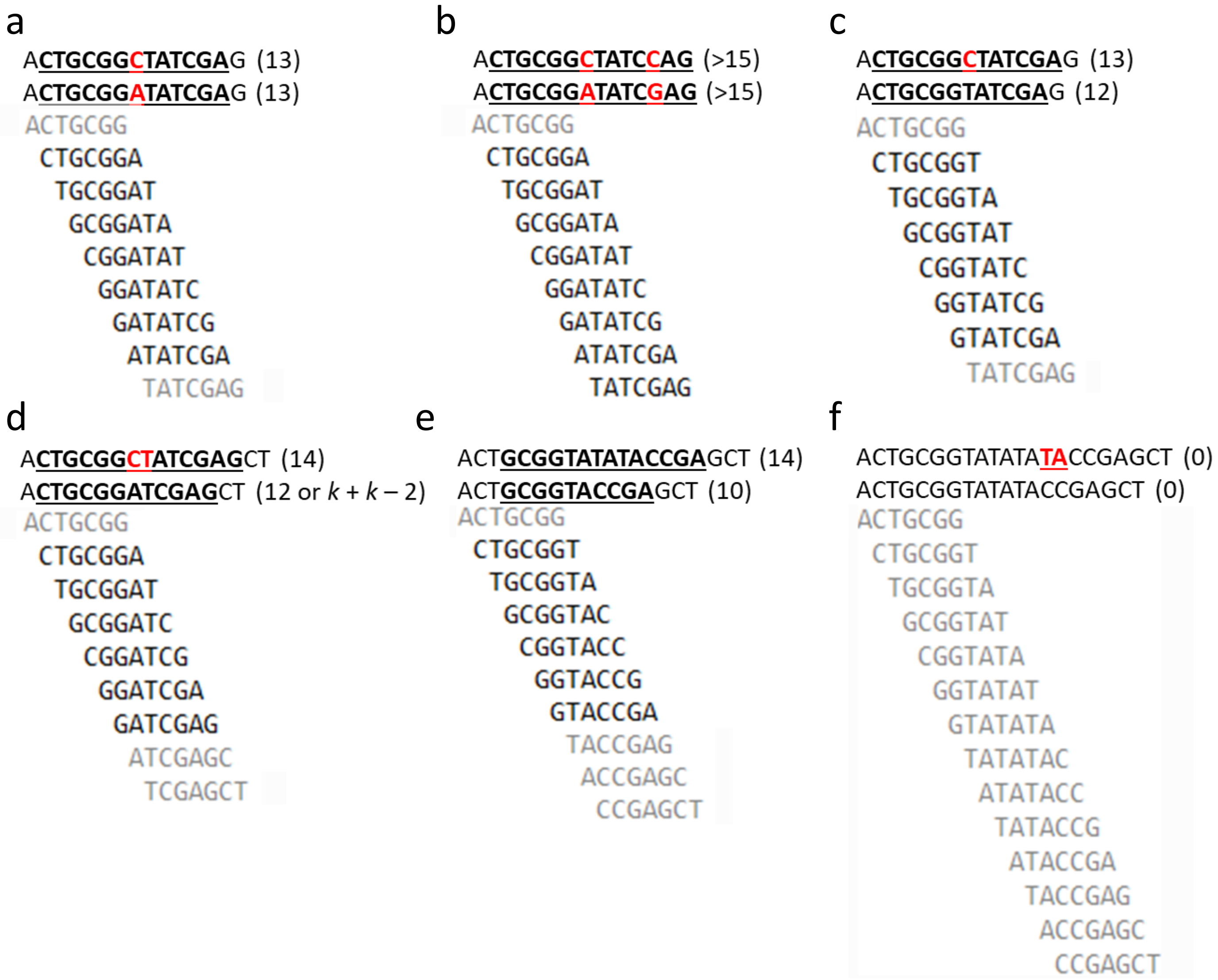


**Figure S4. Associating variants with different fragment sizes.** AFLAP searches for *k*-mers segregating uniquely from either parent. To do this efficiently, *k*-mers are assembled and filtered based on the size of the fragment produced. A single nucleotide variant will produce fragments of $k + k - 1$. For purposes of this illustration $k = 7$, rather than 31 as used in our analysis. Unshaded 7-mers below the sequence are found to be unique to the second sequence. Grey 7-mers are found in both sequences. a) A SNP will result in the retention of 7-mers containing the SNP, 7-mers common to both sequences are filtered out. Using 7-mers the resulting fragment = 13 = $k + k - 1$. b) If two variants are close to one another then larger fragments are assembled because the 7-mers retained contain multiple variants. In this example if no new variants were downstream then the six bases before the C/A variant and six bases after the C/G variant would be retained to assemble a fragment greater than $k + k - 1$. c) Indels may result in fragments equaling less that $k + k - 1$. A single nucleotide deletion results in the six base pairs either side being assembled into a unique 12 bp fragment. 7-mers which contain the nucleotide would still assemble to 13 bp. d) When a dinucleotide deletion is observed a 12 bp fragment is still obtained, 7-mers with the nucleotide now assembled into 14 bp fragments. e) When microsatellite variation is considered much smaller fragments than $k + k - 1$ may be assembled as repetitive sequence leads to the detection of common *k*-mers. f) If the TA repeat was repeated four times and only one copy deleted, then a 7-mer survey would not detect any variants.

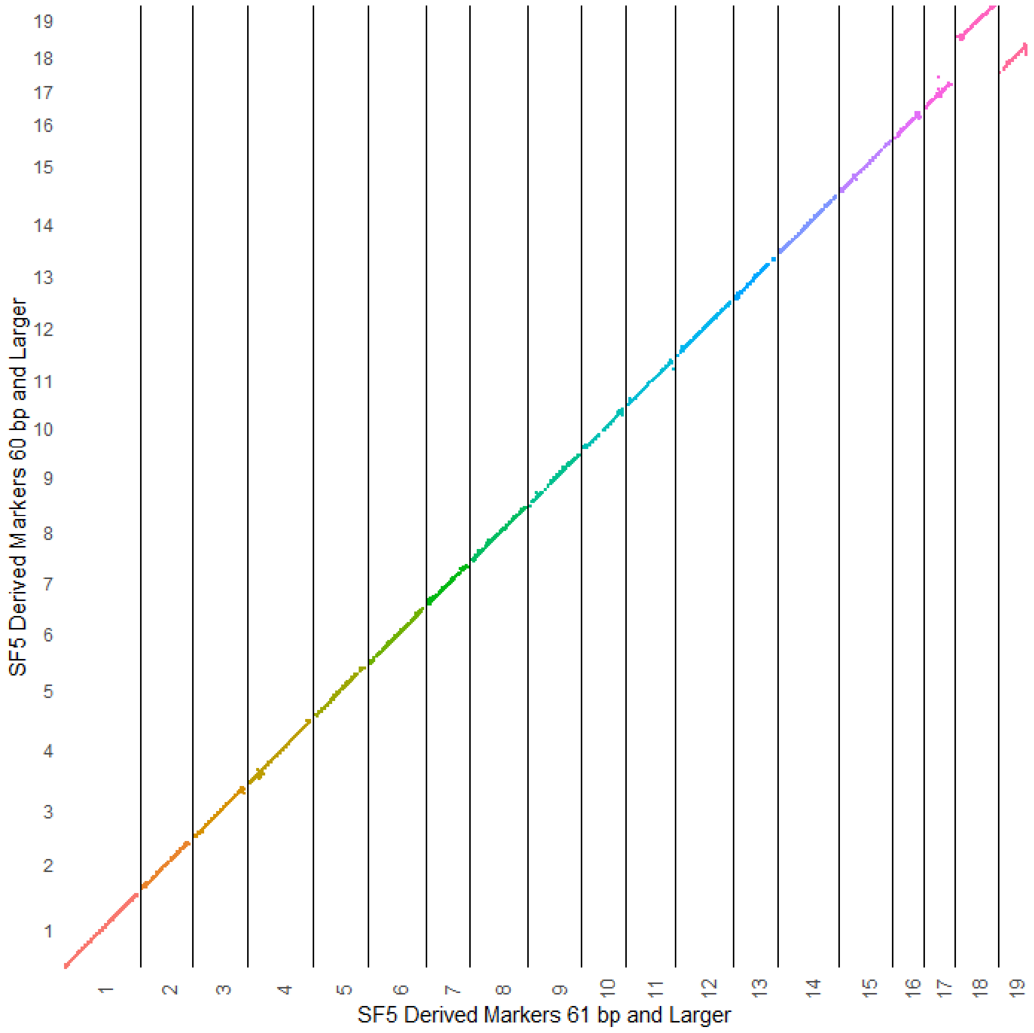


**Figure S5. Addition of markers derived from 60 bp fragments to the SF5 genetic map.** The addition of 4,070 markers derived from 60 bp fragments had little effect on the map as the shared marker order was not altered and the same 19 linkage groups were produced. The switched ordering of linkage group 18 and 19 in the plot is due to more markers derived from 60 bp fragments being assigned to linkage group 19 (1,838 to 1,923) than 18 (1,858 to 1,919).
